# An alphacoronavirus polymerase structure reveals conserved co-factor functions

**DOI:** 10.1101/2023.03.15.532841

**Authors:** Thomas K. Anderson, Peter J. Hoferle, Kenneth W. Lee, Joshua J. Coon, Robert N. Kirchdoerfer

## Abstract

Coronaviruses are a diverse subfamily of viruses containing pathogens of humans and animals. This subfamily of viruses replicates their RNA genomes using a core polymerase complex composed of viral non-structural proteins: nsp7, nsp8 and nsp12. Most of our understanding of coronavirus molecular biology comes from the betacoronaviruses like SARS-CoV and SARS-CoV-2, the latter of which is the causative agent of COVID-19. In contrast, members of the alphacoronavirus genus are relatively understudied despite their importance in human and animal health. Here we have used cryoelectron microscopy to determine the structure of the alphacoronavirus porcine epidemic diarrhea virus (PEDV) core polymerase complex bound to RNA. Our structure shows an unexpected nsp8 stoichiometry in comparison to other published coronavirus polymerase structures. Biochemical analysis shows that the N-terminal extension of one nsp8 is not required for *in vitro* RNA synthesis for alpha and betacoronaviruses as previously hypothesized. Our work shows the importance of studying diverse coronaviruses to reveal aspects of coronavirus replication while also identifying areas of conservation to be targeted by antiviral drugs.

**Significance Statement:** Coronaviruses are important human and animal pathogens with a history of crossing over from animal reservoirs into humans leading to epidemics or pandemics. Betacoronaviruses, such as SARS-CoV and SARS-CoV-2, have been the focus of research efforts in the field of coronaviruses, leaving other genera (alpha, gamma, and delta) understudied. To broaden our understanding, we studied an alphacoronavirus polymerase complex. We solved the first structure of a non-betacoronavirus replication complex, and in doing so identified previously unknown, and conserved, aspects of polymerase cofactor interactions. Our work displays the importance of studying coronaviruses from all genera and provides important insight into coronavirus replication that can be used for antiviral drug development.

## Introduction

The coronavirus (CoV) subfamily is composed of four genera: the alpha, beta, gamma, and delta-CoVs. Across all these genera there are diverse human and animal pathogens that cause a wide range of disease severities [1]. Since 2002, three betaCoVs have emerged from animal reservoirs and caused human epidemics or pandemics: SARS-CoV, MERS-CoV, and SARS-CoV-2 [2–5]. The most recent, SARS-CoV-2, emerged in 2019 and is the causative agent of the COVID-19 pandemic [5, 6]. There are four endemic common-cold causing human CoVs: HuCoV-229E, HuCoV-NL63 and HuCoV-HKU1, HuCoV-OC43 belonging to the alpha or betaCoV genera respectively [1]. In 2021, a recombined feline/canine alphaCoV, named CCoV-HuPn-2018, was identified in samples from human patients with pneumonia in Malaysia and the United States [7–9]. CCoV-HuPn-2018 is the only recorded human emergent alphaCoV in recent years and exemplifies the persistent threat of alphaCoV spillover. In the alphaCoV genera there are several porcine pathogens including transmissible gastroenteritis virus (TGEV), swine enteric alphacoronavirus (SeACoV), and porcine epidemic diarrhea virus (PEDV) [10, 11]. Since 2010, PEDV has had detrimental impacts on the global swine industry, with outbreaks having 70–100% piglet fatality rates at swine farms despite vaccination efforts [12–14].

CoVs are positive-sense, single-stranded RNA viruses with large, ~30 kilobase genomes [1, 15]. To replicate their genomes CoVs encode an RNA-dependent RNA polymerase (RdRP), termed non-structural protein 12 (nsp12) [16–18]. Nsp12 also contains the nidovirus RdRP-associated nucleotidyltransferase (NiRAN) domain at its N terminus, which has been shown to be involved in mRNA capping [19, 20]. *In vitro* studies have shown that viral cofactors nsp7 and nsp8 interact with nsp12 to form an active and processive polymerase complex [21]. The SARS-CoV-2 complex is the fastest known RdRP with nucleotide addition rates up to 170 nt/sec *in vitro* [22]. The first structure of a CoV polymerase complex from the betaCoV, SARS-CoV revealed a subunit stoichiometry of one nsp12, two nsp8, and one nsp7 where nsp7 and one nsp8 form a heterodimer [19]. The nsp8 that directly interacts with nsp12 is denoted as nsp8_1_, and the nsp8 that interacts with nsp7 and nsp12 is denoted as nsp8_2_. In this paper we refer to the complex of nsp12, nsp7, and nsp8 as the CoV core polymerase complex.

Recent SARS-CoV-2 core polymerase structures have provided key insights into betaCoV replication. All SARS-CoV-2 complexes shown to be enzymatically active or bind RNA *in vitro* had the same 1:2:1 nsp7:nsp8:nsp12 stoichiometry as SARS-CoV [23, 24]. Structures determined with longer dsRNA substrates have revealed that each nsp8’s N-terminal extension was bound to upstream dsRNA as it exits the polymerase active site, which has been hypothesized to promote processivity [23, 24]. Addition of the viral RNA helicase, nsp13, to core polymerase complexes showed that each nsp8 subunit independently scaffolds a nsp13 (two nsp13s bind a single core complex) [20, 25, 26]. The nsp13 associated with nsp8_2_ was shown to be capable of binding to the 5’ end of template RNA prior to the RNA entering the polymerase active site [26]. Nsp13’s association with the template RNA produced a conundrum, where nsp12 and nsp13 are oriented such that the polymerase and helicase translocate in opposite directions. Further biochemical and structural work demonstrated that this nsp13 can stimulate template backtracking of the polymerase, during which the 3’ end of nascent RNA extrudes through the nsp12 NTP channel [27]. While this nsp13 has been implicated in backtracking, the triggers and regulation of backtracking are unknown.

To date, there is limited knowledge of CoV polymerase biochemistry and structural biology outside the sarbecovirus subgenus (SARS-CoV and SARS-CoV-2). To our knowledge, *in vitro* demonstration of robust nsp12 polymerase activity has yet to be demonstrated for an alpha-, gamma-, or deltaCoV. Similarly, outside of betaCoVs there is limited structural and biochemical information for nsp7 and nsp8. A crystal structure of the alphaCoV feline coronavirus (FCoV) nsp7-nsp8 complex revealed that alpha- and betaCoV nsp7s and the nsp8 head domains have high structural homology [28]. Across CoV genera there is sequence conservation (>40%) of the nsps involved in RNA synthesis (i.e., nsp7, nsp8, and nsp12) [29]. Sequence and available structural homology suggest shared mechanisms among these viruses to replicate their RNA genomes.

Though our knowledge of SARS-CoV and SARS-CoV-2 replication mechanics has improved over the last two decades, the narrow structural biology focus on two closely related betaCoVs limits our understanding of CoV replication across the virus subfamily. To address this gap, we studied the polymerase core complex of the alphaCoV PEDV using biochemistry and structural biology. Our 3.3 Å structure of the PEDV core polymerase complex bound to an RNA primer-template pair shows the lability of nsp8_2_ to participate in this complex and additional mutagenesis and biochemistry demonstrate that the nsp8_2_ N-terminal helical extension is not necessary for either alpha- or betaCoV RNA synthesis activity. The identification of conserved mechanisms and structural motifs between alpha- and betaCoVs will allow for the development of broadly acting CoV therapeutic strategies.

## Results

### Assembly of an active PEDV polymerase complex

To study the PEDV polymerase complex we recombinantly expressed and purified nsp7, nsp8, and nsp12 (**Fig. S1)**. Assembly of these proteins with a short RNA duplex revealed a complex weight of ~185 kDa (+/- 20 Da) by native mass spectrometry (**Table 1**, **Fig. S2**). This mass is equivalent to one nsp12, one nsp7, two nsp8s, and one RNA duplex. Using an *in vitro* primer extension assay we saw optimal polymerase activity in the presence of all three nsps (**Fig. 1A and Fig. S3**). The necessity of nsp7, nsp8 and nsp12 for robust primer extension is a shared requirement between PEDV, SARS-CoV and SARS-CoV-2 [17, 21]. The PEDV polymerase complex was modestly active (60% of the core polymerase complex) in the absence of nsp7 (**Fig. 1A**). Recent studies showed that MERS-CoV nsp8 and nsp12 can form a modestly active polymerase complex in the absence of nsp7 [30].

**Figure 1:**
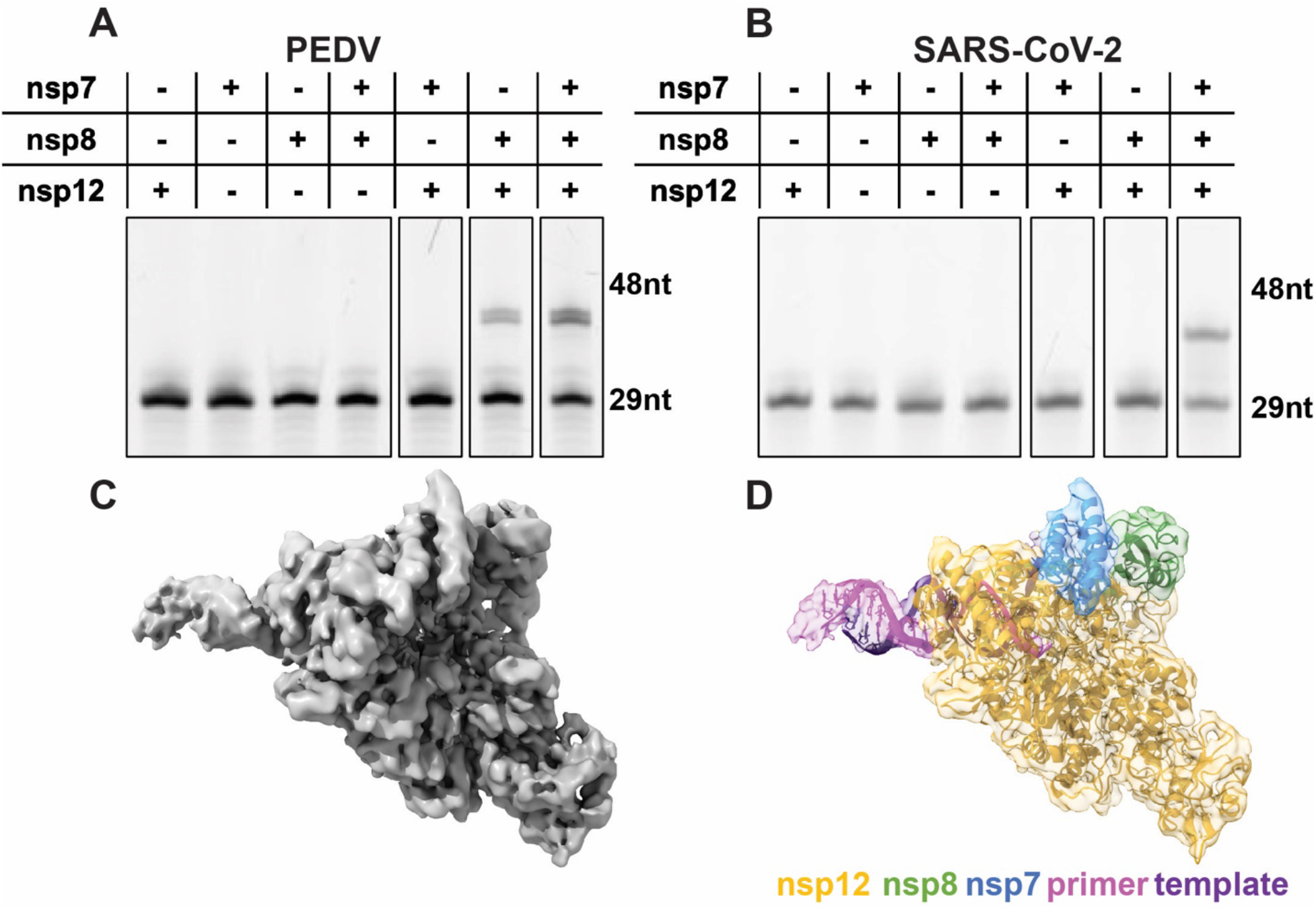
Assembly of an active PEDV polymerase complex. A 29 nt RNA primer with a 5’ fluorophore is annealed to a 38 nt template and extended in the presence of CoV polymerase complexes. Combinations of nsp7, nsp8, and nsp12 were tested for PEDV (**A**) and SARS-CoV-2 (**B**). **C**) 3.4 Å cryo-EM reconstruction of the PEDV core polymerase complex. **D**) Coordinate model of the PEDV core polymerase complex docked into its corresponding electron density map colored by chain.

**Table 1:** Native mass spectrometry of coronavirus polymerase complexes. Major and minor species from native mass spectrometry experiments are listed for each complex tested with their respective (±) standard deviations. The expected protein or complex associated with each weight are as follows: ~22 kDa – nsp8, ~110 kDa – nsp12, ~185 kDa – intact core complex, 218 kDa – nsp12 dimer, 370 kDa – complex dimer (See also **Table S2** and **Figure S4**).

| Sample | Major Peak(s)<br>(Da) | Minor Peak(s)<br>(Da) |
| --- | --- | --- |
| PEDV nsp12+8+7 | 185,120 ± 70 | 109,450 ± 40 |
| PEDV A382R-nsp12+8+7 | 109,533 ± 60<br>185,120 ± 30 | 218,880 ± 50 |
| PEDV V848R-nsp12+8+7 | 185,200 ± 200 | 370,700 ± 600<br>109,630 ± 200 |
| PEDV<br>nsp12+8+8L7 | 184,670 ± 60 | 109,420 ± 30 |
| SARS-CoV-2<br>nsp12+8+7 | 186,240 ± 70 | 110,680 ± 20 |
| SARS-CoV-2<br>L387R-nsp12+8+7 | 186,400 ± 100 | 110,900 ± 100<br>22,400 ± 100 |
| SARS-CoV-2<br>T853R-nsp12+8+7 | 186,190 ± 50 | 22,300 ± 50 |
| SARS-CoV-2<br>nsp12+8+8L7 | 186,270 ± 40 | 110,600 ± 40 |

### Structure of PEDV core polymerase complex

Using single-particle cryo-EM we determined the structure of the PEDV core polymerase complex (nsp7, nsp8, nsp12) bound to a short RNA primer-template pair in a posttranslocated state (**Fig. 1C and D**). Our final map has a resolution of 3.3 Å (**Table S1, Fig S4**). Our cryo-EM map contains density for nsp12 residues 3-355, 362-890, and 907-923, which includes the NiRAN domain and polymerase domains. We resolved most of nsp7 (residues 2-62), the C-terminal portion of nsp8_1_ (residues 79-192), and a full turn of dsRNA exiting the polymerase active site. We lack well-resolved density for the nsp8_1_ N-terminal helical extension. 3D variability analysis showed that the dsRNA leaving the active site and N-terminal extension of nsp8_1_ have flexibility, likely leading to their lack of density in the final reconstruction (**Video S1**) [31]. Unexpectedly, our final reconstruction lacked any density for nsp8_2_, differing with most published CoV complex structures as well as the PEDV polymerase native mass spectrometry data described above.

### Comparison of PEDV and SARS-CoV-2 polymerase core complex models

PEDV and SARS-CoV-2 nsp7, nsp8, and nsp12 protein sequences have significant sequence identity (nsp12 58.6%, nsp8 43.1%, and nsp7 41.8%) (**Fig. S5**). The overall architecture of the PEDV polymerase core complex is very similar to published SARS-CoV-2 models with a nsp12 RMSD value of 0.964 Å [27]. Comparison of our PEDV model’s RdRP active site with SARS-CoV-2 structures revealed highly conserved active site structures (**Fig. S6**). NiRAN architectures between PEDV and SARS-CoV-2 nsp12 are also very similar. This structural conservation suggests that antivirals targeting SARS-CoV-2, such as Remdesivir or the recently studied dual-purpose AT-527, could be effective against PEDV and alphaCoVs [32–34].

A major difference in the PEDV model is an altered loop conformation (PEDV nsp12 residues 249-268) that binds the nsp8_1_ head domain in PEDV but not in SARS-CoV-2 (**Fig. 2A**). This conformational difference increases nsp8_1_’s buried surface area by 115 Å^2^ in the PEDV polymerase core complex. This PEDV nsp12 region is highly conserved within alphaCoV but not across CoV genera (**Fig. S5**). From these observations we predict that the increased nsp8_1_ buried surface area is shared among and specific to alphaCoVs.

**Figure 2:**
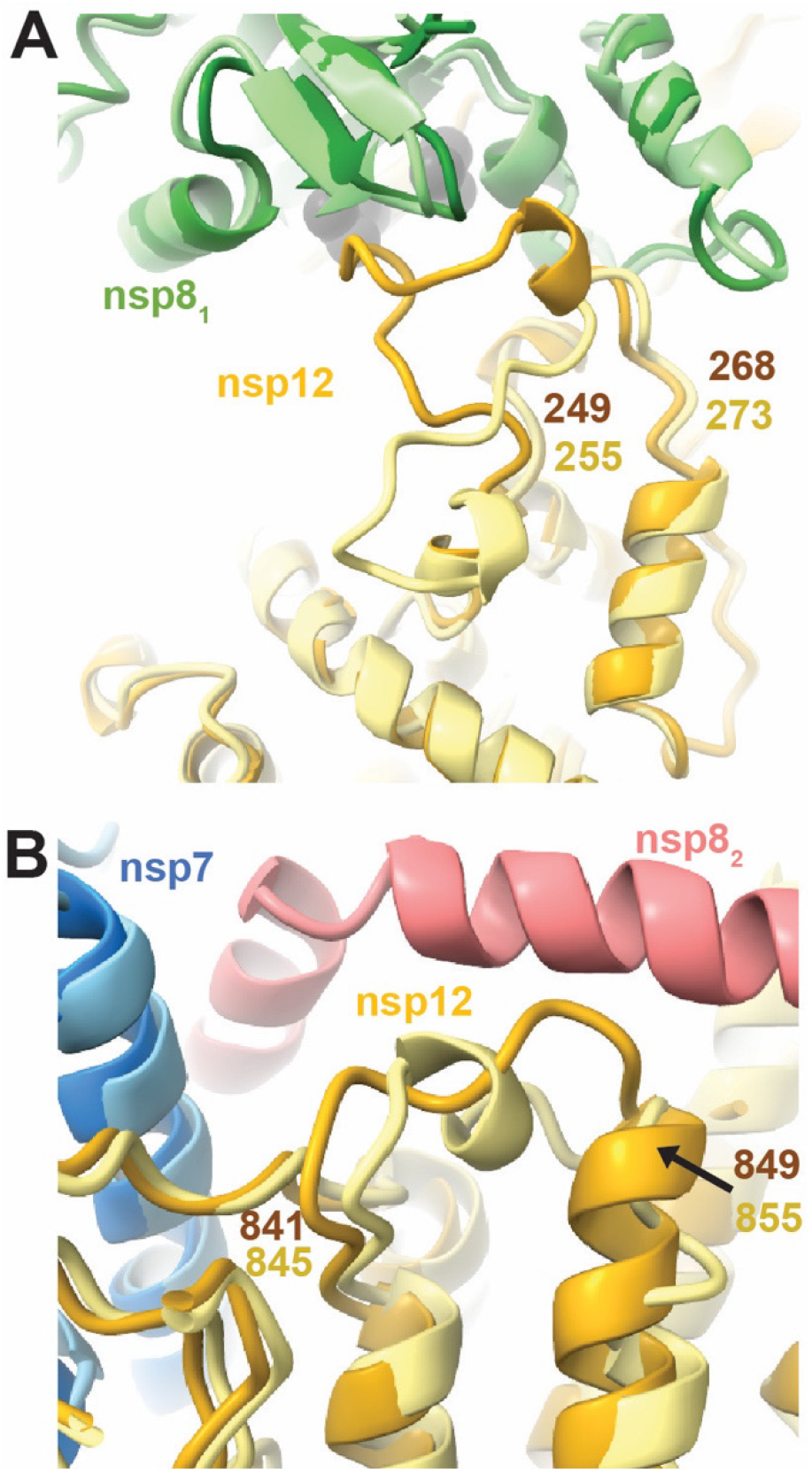
Structural differences between PEDV and SARS-CoV-2 nsp12s. **A**) PEDV nsp12 loop (residues 249-268) buries extra surface area on nsp8_1_ compared to SARS-CoV-2 (PDB: 7KRP). **B)** PEDV nsp12 841-849 lacks helical definition in the absence of nsp8_2_, while SARS-CoV-2 nsp12 845-855 (PDB: 7KRP) has a short helix when the nsp8_2_ N-terminal extension is present. For each panel PEDV and SARS-CoV-2 models are superimposed with PEDV being colored in darker shades. Note that nsp8_2_ is not observed in the PEDV structure.

The PEDV nsp12 residues 841-849 lack a helical definition (**Fig. 2B**) observed in SARS-CoV-2 models with resolved nsp8_2_ N-terminal helical extensions (PDB IDs: 6YYT, 7KRN, 7KRO, 6XEZ). However, SARS-CoV (PDB ID: 6NUR) and SARS-CoV-2 structures (PDB IDs: 7CYQ, 7CXM) without well resolved nsp8_2_ extensions also lack helical definition in this region suggesting that association of the nsp8_2_ N-terminal helical region promotes helical secondary structure in this nsp12 region.

### PEDV nsp12 structure model has an alternate binding stoichiometry with nsp8

Missing density for nsp8_2_ in our reconstructed map was unexpected. Prior structural work for betaCoV polymerases has proposed that nps82 participates in CoV RNA synthesis, acting with nsp8_1_ as “sliding-poles” that guide dsRNA out of the polymerase active site [23]. Our native mass spectrometry results also contrast the PEDV polymerase structural model’s stoichiometry (**Table 1**). These observations led us to hypothesize that nsp8_2_’s interactions with the polymerase complex are weaker than other cofactors (nsp8_1_ and nsp7), and perhaps the vitrification process during grid freezing caused nsp8_2_ disassociation. Interestingly, the PEDV complex remained bound to RNA in the absence of nsp8_2_. We therefore hypothesized that nsp8_2_ may not be required for CoV RNA binding or RNA synthesis.

*In vitro* primer extension demonstrates that for both PEDV and SARS-CoV-2 proteins, optimal polymerase activity is achieved in the presence of all three nsps, but there is modest activity in the absence of nsp7 for PEDV (60% of wildtype) (**Fig. 1A and B**). As the association of nsp8_2_ with the core complex is largely through binding to nsp7, modest polymerase activity in the absence of nsp7 suggests a strong role for nsp8_1_ in stimulating core complex polymerase activity. This is similar to a previous study of MERS-CoV polymerase where nsp7 and hence the nsp7-nsp8_2_ heterodimer was not required for in vitro RNA synthesis [30]. To dissect the different nsp8s’ contributions to promoting polymerase activity, we designed PEDV and SARS-CoV-2 nsp12 point mutations at each nsp8’s proteinprotein interfaces. These mutations were designed to be highly disruptive to hydrophobic protein-protein interfaces through the substitution of nsp12 surface residues with arginine to prevent association of a nsp8 (**Fig. 3A and B**). The effects of these mutations were examined using the *in vitro* primer extension assay (**Fig. 3C, D and Fig. S7**).

**Figure 3:**
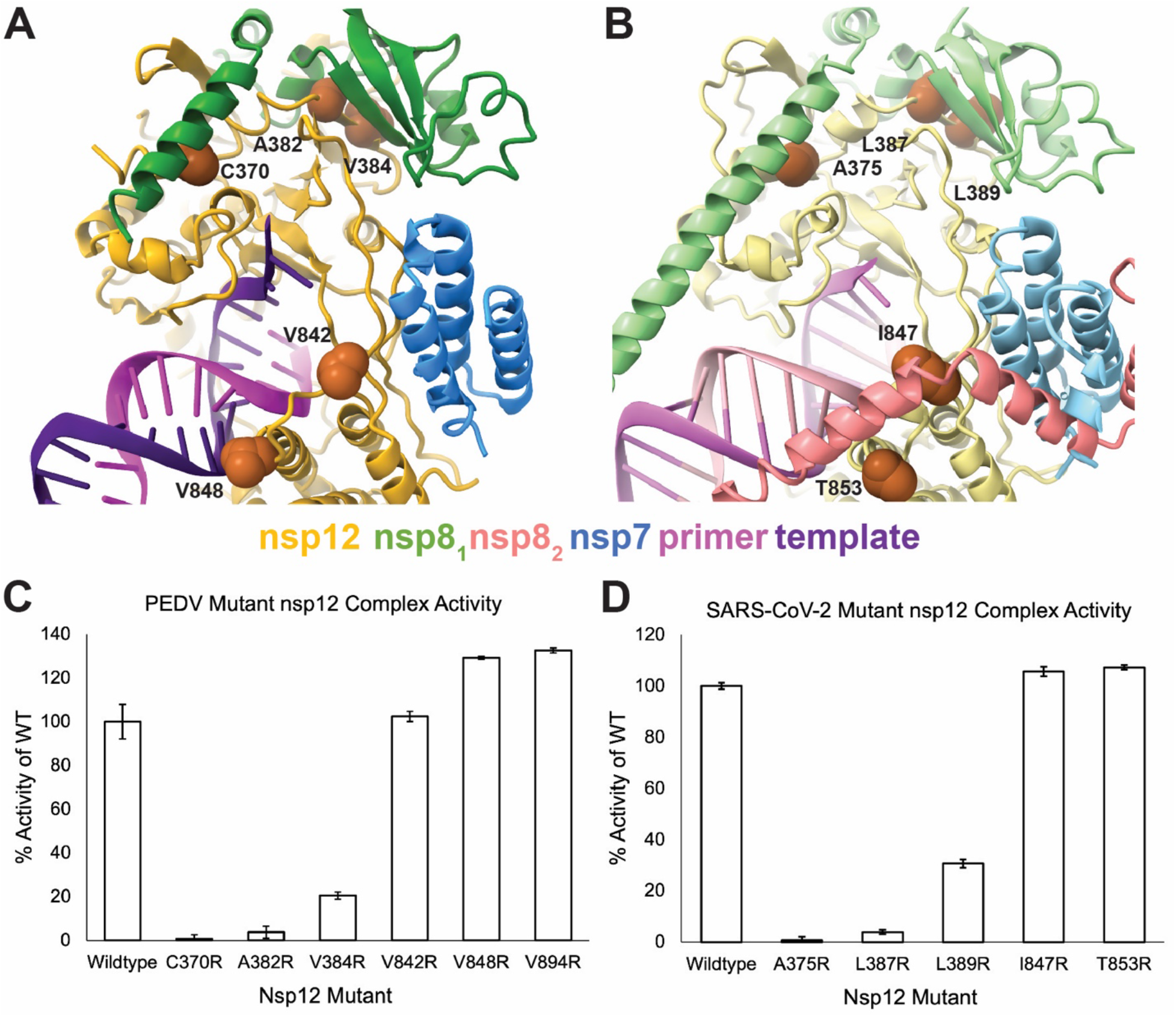
Effects of cofactor disrupting nsp12 mutations on polymerase activity. Sites of nsp12 mutations designed to disrupt cofactor interactions for **(A)** PEDV and **(B)** SARS-CoV-2 are shown as orange spheres. PEDV nsp12 residue V894 is not indicated due to lack of density. The effects of PEDV (**C**) and SARS-CoV-2 (**D**) nsp12 mutations were evaluated using *in vitro* RNA primer extension assays. Technical triplicates of reactions were run and percent activity of wildtype nsp12 is presented. Error bars indicate standard deviation of the triplicates. All reactions, except PEDV nsp12-V842R, were significantly (P<0.05) different compared to the wildtype control. Significance was determined using an unpaired t-test for each mutant complex.

Nsp12 mutants designed to disrupt the association of PEDV nsp8_1_ N-terminal helical extension, (nsp12 C370R) or nsp8_1_ C-terminal head domain (nsp12 A382R or V384R) both resulted in complexes that either completely or mostly lost RNA extension activity (**Fig. 3C and Fig. S7**). In contrast, mutant complexes designed to disrupt association of nsp8_2_ N-terminal helical extension (nsp12 V842R, V848R, or V849R), which were based on structural homology with the SARS-CoV-2 core complex (PDB ID: 6YYT), retained polymerase activity (**Fig. 3C**). These results indicate the importance of nsp8_1_ for PEDV RNA synthesis and demonstrate that the nsp8_2_ N-terminal helical extension is not required for stimulation of polymerase activity.

To test if these findings are specific to the PEDV polymerase core complex or a shared feature among CoVs, we produced and tested homologous mutations for SARS-CoV-2 nsp12. Like PEDV, SARS-CoV-2 nsp12 mutants designed to disrupt association of the nsp8_1_ N-terminal extension (nsp12 A375R) or nsp8_1_ C-terminal head domain (nsp12 L387R or L389R) resulted in inactive polymerase complexes (**Fig. 3D and Fig. S8**). However, nsp12 mutations designed to disrupt association of the nsp8_2_ N-terminal extension (I847R or T853R) produced polymerase complexes with activity resembling that of the wild-type protein (**Fig. 3D**). Similar results for mutagenesis across both PEDV and SARS-CoV-2 core polymerase complexes suggest shared roles for nsp8s across the coronavirus subfamily.

Native mass spectrometry (**Table 1**, **Fig. S3**) of select mutant complexes showed full complex stoichiometry, 1:2:1:1 nsp7:nsp8:nsp12:RNA (~186 kDa mass), despite the nsp12 mutations. The only exception being PEDV A382R, where there were two major species of solo nsp12 and intact core polymerase complexes, indicating that nsp12 was either completely free or completely bound by nsp7 and nsp8 cofactors. These data suggest that the disruptive mutations are only effective in blocking the nsp12-nsp8 subdomain interactions and are not sufficient to fully prevent nsp8 subunit association with nsp12. This is congruent with the extended nature of nsp8 and the presence of both an N-terminal helical extension and a C-terminal head domain that bind distinct sites in coronavirus core polymerase complexes for both nsp8_1_ and nsp8_2_. Hence nsp12 mutations disrupting the binding of one region of nsp8 may not preclude association of that nsp8 with the core complex. These data also suggest that both the nsp8_1_ N-terminal extension and the C-terminal head domain play essential roles in stimulating polymerase activity beyond simply facilitating protein-protein interactions.

### Nsp8_2_ N-terminal RNA binding domain is not required for robust RNA synthesis activity

Recent work showed that a nsp8-nsp7 (nsp8L7) fusion protein can efficiently function as a nsp12 cofactor and allows the contributions of each nsp8 of a coronavirus core complex polymerase activity to be delineated [35]. We produced SARS-CoV-2 and PEDV nsp8L7 fusion proteins with six-residue linker regions (**Fig. 4A**, **Fig. S1**). Both SARS-CoV-2 and PEDV nsp8L7 fusion proteins could stimulate their respective polymerase core complex activities in the presence of free nsp8 (**Fig. 4B and C**). While SARS-CoV-2 nsp8L7 resulted in activity similar to wildtype nsp7 and nsp8, PEDV nsp8L7 only resulted in 60% stimulation compared to the use of wildtype. It has previously been shown that nsp8 alone, and nsp7 and 8 together form different oligomeric states depending on the CoV genera [36]. The PEDV polymerase activity with nsp8L7 is similar to PEDV reactions lacking nsp7 and we predict that the reduction in activity (compared to wildtype) is due to a reduced dissociation of the fusion construct from oligomeric states into forms available for binding nsp12. Therefore, only a fraction of the available nsp12 may form full complexes as compared to SARS-CoV-2 nsp8L7 complexes (**Table 1**, **Fig S2**). In support of nsp8_1_ being required for RNA synthesis *in vitro* and in validation of the fusion protein strategy to delineate nsp8 contributions, the nsp8L7 fusion protein is not sufficient to stimulate CoV RNA synthesis in the absence of isolated nsp8 (**Fig 4B and C**).

**Figure 4:**
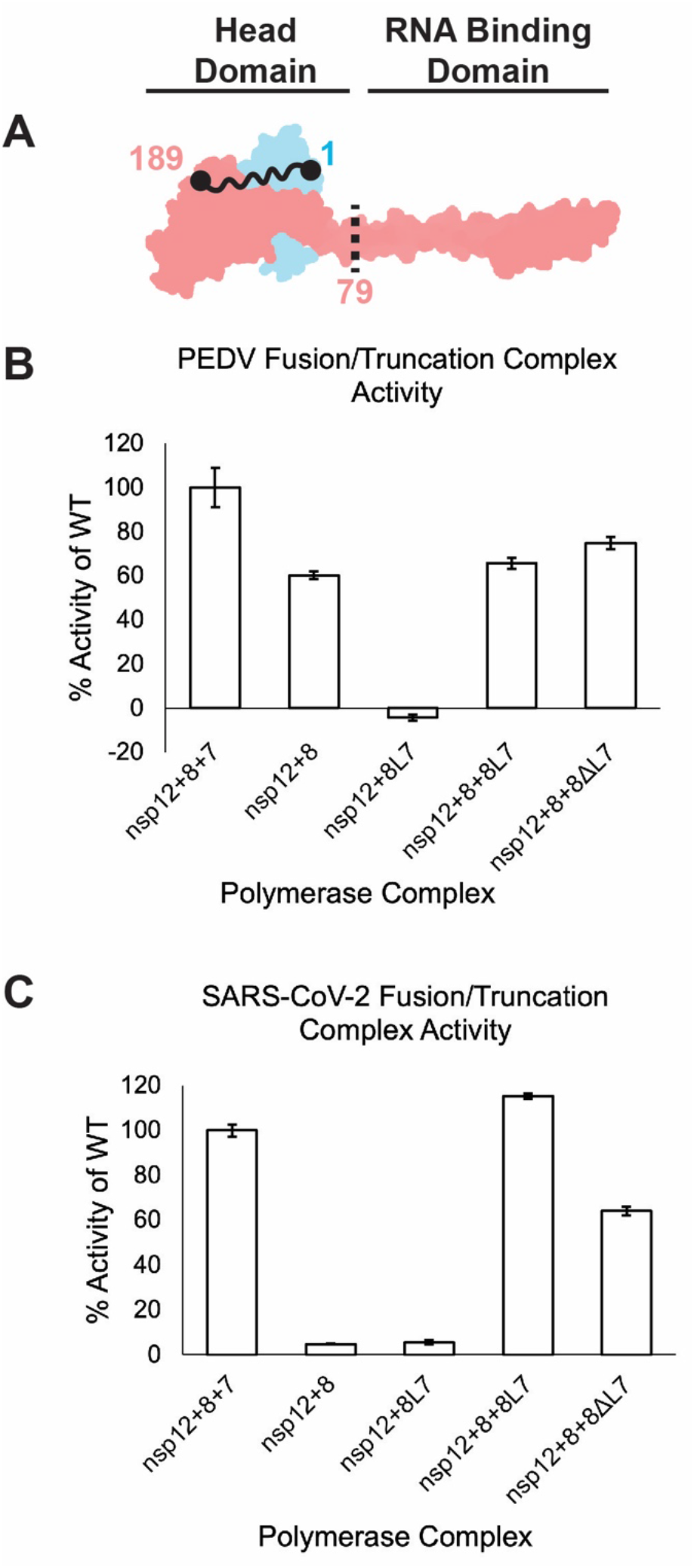
nsp8_2_’s N-terminal domain is not required for RNA synthesis. **A)** Model of nsp7 (blue) and nsp8 (red) heterodimer (PDB ID: 7KRP). The black dots and squiggly line depict the nsp8 and nsp7 termini that are fused in nsp8L7. The dashed black line is the site of truncation of nsp8ΔL7, removing the nsp8_2_ N-terminal 79 amino acid RNA binding domain. **B)** PEDV and **C)** SARS-CoV-2 polymerase complex activity using the nsp8L7 and nsp8ΔL7 co-factors. Reactions were run in triplicate and percent activity was compared to a wildtype nsp7+nsp8+nsp12 complex. Error bars indicate standard deviation of the triplicates. Each reaction was determined to be significantly different than all other complexes tested (p<0.05) using an unpaired t-test.

Using the PEDV and SARS-CoV-2 nsp8L7 fusion constructs, we produced 79 residue N-terminal truncations of nsp8_2_ (nsp8ΔL7) which lack the previously described RNA binding region of the protein (**Fig. 4A, Fig. S1)** [23]. Both PEDV and SARS-CoV-2 nsp8ΔL7 stimulated core polymerase primer extension activity *in vitro* (**Fig. 4**). Compared to the activity of the fusion constructs, PEDV had a 20% increase and SARS-CoV-2 had a 60% decrease. PEDV’s ~20% increase in activity (80% of wildtype) compared to reactions with the fusion construct, or those lacking nsp7 indicate that the nsp8_2_ N-terminal region is not required for RNA synthesis *in vitro.* SARS-CoV-2’s 60% activity (compared to wildtype) is a large increase in activity over the non-active nsp8 + nsp12 reactions, showing that nsp8ΔL7 acts as a functional cofactor, albeit not as well as nsp8L7. Once again, the differences in PEDV and SARS-CoV-2 activities could be attributed to the cofactor proteins unique interactions across the CoV genera [37]. While the effects of the PEDV and SARS-CoV-2 truncations vary, the fact that both truncation reactions have increased activity compared to reactions lacking nsp7 indicates that nsp8ΔL7 is a functional cofactor and that the N-terminal extension of nsp8_2_ is not required for alpha- or betaCoV RNA synthesis *in vitro*.

## Discussion

In this study, we establish that the PEDV core polymerase complex assembles into an active polymerase complex using a similar cofactor stoichiometry as was previously seen in betaCoV. Our structure of the PEDV core polymerase complex reveals an overall similar architecture to betaCoV with a large conformational difference in PEDV nsp12 residues 249-269 to interact with nsp8_1_ that we predict to be conserved among alphaCoV. Additionally, we used biochemistry and mutagenesis to show that for both PEDV and SARS-CoV-2, nsp8_1_ is required for RNA synthesis while the N-terminal extension of nsp8_2_ is not.

Since observed structural differences in the PEDV and SARS-CoV-2 core polymerase complexes are outside of the common targets for antiviral drug design (the polymerase and NiRAN active sites), we predict that antivirals targeting these shared sites would be effective against both alpha- and betaCoV. For example, structures have shown that SARS-CoV-2 nsp12 residue S861 is likely important for the effectiveness of the nucleotide analogue Remdesivir [38]. The predicted mechanism is that S861 sterically classes with the 1’-cyano group of the antiviral, impairing RNA elongation. For PEDV, the residue S856 is conserved in both sequence and space to S861 of SARS-CoV-2 (**Fig. S6**), indicating that Remdesivir could be used to treat alphaCoV infections [39].

Although our structural model of the PEDV core polymerase complex lacks nsp8_2_, native mass spectrometry confirmed nsp8_2_’s association with the complex. *In vitro* analyses with mutant PEDV and SARS-CoV-2 nsp12s and truncated nsp8_2_ (nsp8L7) established that nsp8_2_’s N-terminal extension is not required for *in vitro* RNA synthesis. Previous work has suggested that the N-terminal extensions of nsp8_1_ and nsp8_2_ act as sliding poles to promote RNA synthesis and processivity [23]. We propose a revision to this model where the nsp8_2_ N-terminal extension is not essential for RNA synthesis but rather may have an important role in other stages of replication.

We hypothesize that nsp8_2_ may act as a scaffolding protein that regulates the association of other viral factors like the nsp13 RNA helicase to promote RNA backtracking (**Fig. 5**). Previous work has shown that the viral helicase, nsp13, binds template RNA entering the active site and can cause polymerase backtracking on primer-template RNA [27]. In a backtracked polymerase state, the 3’ end of the nascent RNA extrudes out of the polymerase NTP entry channel. While a function for backtracking within viral replication remains unclear, one hypothesis is that it mediates proofreading of mis-incorporated 3’ nucleotides during CoV RNA synthesis. The nsp13 responsible for backtracking RNA binds the polymerase core complex via the nsp8_2_ N-terminal extension. As a potential model for viral RNA proofreading, we propose that aberrant nucleotide incorporation stalls elongation, allows association of the nsp8_2_ N-terminal extension and the backtracking nsp13 RNA helicase with the polymerase core complex to induce a backtracked state. 3’ nucleotides of the backtracked RNA could then be excised by the viral nsp14 exonuclease removing the misincorporated nucleotides.

**Figure 5:**
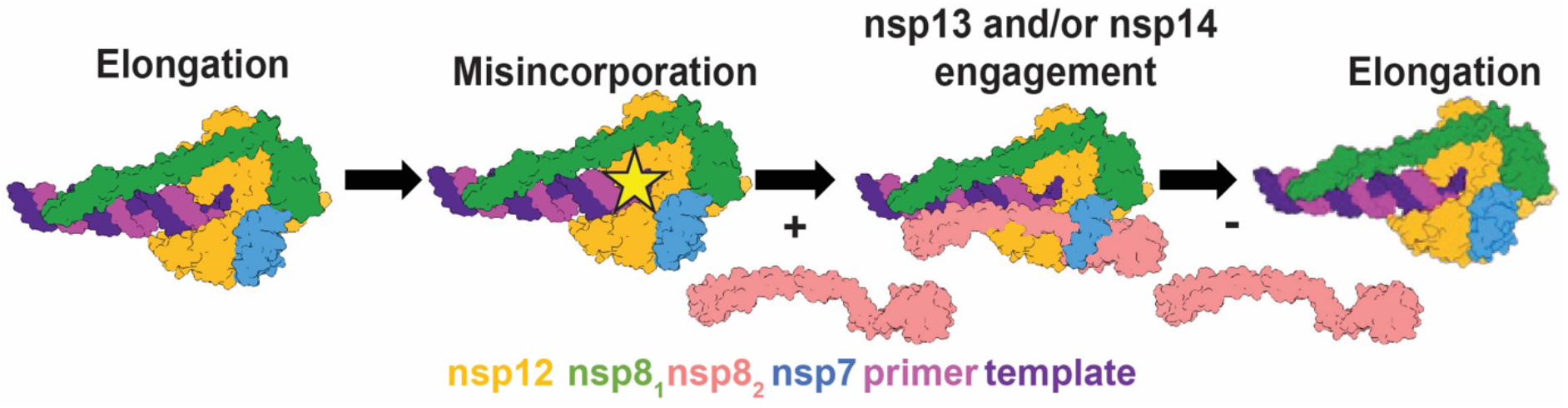
Model for coronavirus backtracking and proofreading. Stalling of polymerase complexes by misincorporation is hypothesized to promote the engagement of nsp8_2_’s N-terminal helical extension with RNA allowing the recruitment of additional viral factors for backtracking or proofreading. Whether or not nsp8_2_ is always associated with the complex is unknown. Here, for simplicity, we depict the polymerase core elongation complex without nsp8_2_.

This work expands the diversity of CoV polymerase structural biology to the alphaCoVs. Our work shows a high level of structural conservation among CoV replication machinery and has allowed for the generation of hypotheses for CoV RNA synthesis that span more than a single virus genus. Such work is essential not only for a greater understanding of CoV biology but also in preparation for the threat of emerging CoV.

## Materials and Methods

### Expression Construct Design

DNA encoding PEDV and SARS-CoV-2 nsps were codon optimized and synthesized (Genscript). PEDV protein sequences correspond to GenBank AKJ21892.1. SARS-CoV-2 protein sequences correspond to GenBank UHD90671.1. PEDV nsp7 and nsp8 genes were cloned into pET46 and pET45b expression vectors, respectively. PEDV nsp7 was cloned with a C-terminal TEV protease site and hexahistidine tag. PEDV nsp8 has an N-terminal hexahistidine tag and TEV protease site. SARS-CoV-2 nsp7 and nsp8 were both cloned into pET46 with N-terminal hexahistidine tags, and enterokinase and TEV protease sites. Both PEDV and SARS-CoV-2 nsp12 genes were cloned into pFastBac with C-terminal TEV protease sites and double Strep II tags.

Expression plasmids for nsp12 point mutants were produced using mutagenesis on the pFastBac plasmids. The nsp8-nsp7 (nsp8L7) fusion constructs were produced by overlap PCR creating a GSGSGS peptide linker between nsp8 and nsp7 and inserted into pET46 expression vectors with N-terminal hexahistidine tags, and enterokinase and TEV protease sites. Truncation constructs were produced using Kinase-Ligase-DpnI (KLD) cloning. All open reading frames on the DNA plasmids were verified by Sanger sequencing.

### Recombinant Protein Expression

Recombinant nsp7 and nsp8: Co-factor proteins were expressed in Rosetta 2pLysS *E. coli* cells (Novagen). Cultures were grown at 37°C and induced at an OD_600_ of 0.6-0.8 with isopropyl β-D-1-thiogalactopyranoside (IPTG) at a final concentration of 500 μM. After growing for 16 hours at 16°C, cells were harvested by centrifugation and resuspended in co-factor wash buffer (10 mM Tris-Cl pH 8.0, 300 mM sodium chloride, 30 mM imidazole, and 2 mM DTT). Cells were lysed in a microfluidizer (Microfluidics) and lysates cleared via centrifugation. Co-factors were purified using Ni-NTA agarose beads (Qiagen), eluting with 300 mM imidazole. Eluted protein was digested with 1% (w/w) TEV protease overnight while dialyzing (10 mM Tris-Cl pH 8.0, 300 mM sodium chloride, and 2 mM DTT) at 4°C. Digested protein was flowed back over Ni-NTA agarose beads to removed undigested protein, and further purified using Superdex 200 Increase 10/300 GL (Cytiva) in 25 mM Tris-Cl pH 8.0, 300 mM sodium chloride, and 2 mM DTT. Fractions containing the protein of interest were concentrated using ultrafiltration. Concentrated protein was aliquoted, flash frozen in liquid nitrogen, and stored at −80°C. Co-factor protein yields for 1 L of cells ranged from 10-40 mg.

Recombinant nsp12s: pFastBac plasmids carrying the nsp12 gene and DH10Bac *E. coli* (Life Technologies) were used to produce recombinant bacmids for each gene. Bacmids were transfected into Sf9 cells (Expression Systems) with Cellfectin II (Life Technologies) to produce recombinant baculoviruses, which were twice amplified using Sf9 cells. Amplified baculoviruses were used to infect Sf21 cells (Expression Systems) for protein expression. After two days of incubation at 27°C, cells were collected via centrifugation and pellets resuspended in wash buffer (25 mM HEPES pH 7.4, 300 mM sodium chloride, 1 mM magnesium chloride, and 5 mM DTT) with an added 143 μL of BioLock (IBA) per 1 L of culture. Cells were lysed using a microfluidizer (Microfluidics) and lysates cleared via centrifugation. Protein was affinity purified using Streptactin superflow agarose (IBA) and eluted with wash buffer that contained 2.5 mM desthiobiotin. Protein was further purified via size exclusion chromatography with a Superdex 200 Increase 10/300 GL column (Cytiva) in 25 mM HEPES pH 7.4, 300 mM sodium chloride, 100 μM magnesium chloride, and 2 mM TCEP. Fractions containing nsp12 were pooled and concentrated using ultrafiltration. Concentrated protein was aliquoted, flash frozen in liquid nitrogen and stored at −80°C. Average protein yield for 1 L of culture was 3-5 mg.

### Preparation of RNA Substrates

RNA oligos were purchased from Integrated DNA Technologies. Primer RNA was modified with a 5’ fluorescein to monitor the RNA by gel electrophoresis. Primer RNA: CAUUCUCCUAAGAAGCUAUUAAAAUCACA. Template RNA: AAAAAGGGUUGUGAUUUUAAUAGCUUCUUAGGAGAAUG. RNA template was always held in slight excess of primer (primer: template ratio of 1:1.2). RNA oligos were annealed in 2.5 mM HEPES pH 7.4, 2.5 mM potassium chloride, and 0.5 mM magnesium chloride and heated at 95°C for 5 minutes, then allowed to cool slowly back to 25°C for 75 minutes before either being used immediately or stored at −20°C.

### Native Mass Spectrometry

Component concentrations were, unless stated otherwise, 10.4 μM nsp12, 20.8 μM nsp7, 31.2 μM nsp8 (or nsp8-nsp7 fusion), and 12.5 μM RNA duplex. Proteins were combined in native-MS buffer (10 mM Tris-Cl pH 8.0, 100 mM ammonium acetate, 2 mM magnesium chloride, and 1 mM DTT) and incubated at 25°C for 15 minutes. RNA duplex was then added, and reactions were incubated at 25°C for an additional 15 minutes. SARS-CoV-2 T853R and PEDV V848R complexes (homologous mutations) were prepared at half the normal protein and RNA concentrations due to low yield of the mutant nsp12s.

All samples were buffer exchanged into 100 mM ammonium acetate (NH_4_OAc) with 2 mM MgCl_2_. First, 350 μL 100 mM NH_4_OAc with 2 mM MgCl_2_ was added to a 100 kDa molecular weight cut-off Amicon Ultra-0.5 Centrifugal Filter Unit (Millipore Sigma), followed by 15 μL of ~10 μM polymerase sample. The samples were centrifuged at 14,000 x g for 10 min to load the sample, followed by two rinses with 400 μL of the NH_4_OAc/MgCl_2_ solution, also centrifuged at 14,000 x g for 10 min. Finally, the filter was inverted into a new catch tube and centrifuged at 1,000 x g for 2 min to recover the filtered and concentrated sample. Samples were diluted using the same NH_4_OAc/MgCl_2_ solution to roughly 1–5 μM for introduction to the mass spectrometer via electrospray ionization.

Standard wall borosilicate tubing (1.20 mm o.d., 0.69 mm i.d., Sutter Instrument) was pulled using a P-1000 Micropipette Puller (Sutter Instrument) to a tapered tip of ~3–5 μm diameter. Each sample was loaded into a pulled glass capillary and 1.1–1.3 kV was applied to the sample using a platinum wire inserted into the back of the capillary. Charged droplets entered a Q Exactive UHMR Hybrid Quadrupole Orbitrap Mass Spectrometer (ThermoFisher Scientific) via a heated inlet capillary. The heated capillary was set to 200–250°C to minimize solvent adduction to analyte ions. Additional removal of adducts was accomplished using in-source trapping with a range of injection voltages typically between 150–300 V. The voltage was tuned for each sample to maximize adduct removal with minimal dissociation of the ionized complex. All reported spectra are an average of 50 scans collected with the Orbitrap mass analyzer at a resolution setting of 6,250. Charge states were assigned manually by minimizing the standard deviation of masses calculated by peaks within a single charge state distribution. The average mass and standard deviation are reported for major distributions in each spectrum.

### In vitro Primer Extension Assay

Assay conditions were 10 mM Tris-Cl pH 8.0, 10 mM sodium chloride, 2 mM magnesium chloride, and 1 mM DTT with a typical reaction volume of 20 μL. Protein final concentrations were 500 nM nsp12, 1.5 μM nsp7, and 1.5 μM nsp8 (nsp8-nsp7 fusion and truncation proteins were also at 1.5 μM). Duplex RNA final concentration was 250 nM. Prior to use, proteins were diluted in assay buffer. Diluted proteins were then combined and incubated at 25°C for 15 minutes, duplex RNA was added and reactions incubated at 25°C for an additional 15 minutes. Reactions were initiated by the addition of NTPs to a final concentration of 40 μM and reactions ran for 1 minute (at 25°C for SARS-CoV-2 polymerases, or 30°C for PEDV polymerases) before being halted by addition of two volumes of sample loading buffer (95% (v/v) formamide, 2 mM EDTA, and 0.75 mM bromophenol blue). Samples were heated at 95°C then analyzed using denaturing urea-PAGE (8 M urea, 15% polyacrylamide) run in 1X TBE (89 mM Tris-Cl pH 8.3, 89 mM boric acid, 2 mM EDTA). Gels were imaged using a Typhoon FLA 9000 (GE Healthcare) to identify fluorescein signals. Extension was quantified using ImageJ [40].

### Sample Preparation and Grid Freezing for CryoEM

PEDV polymerase complexes were prepared at a total protein concentration of 1 mg/mL with a ratio of 2: 2: 1: 1.2 nsp7: nsp8: nsp12: RNA duplex. The complex was assembled in 25 mM HEPES pH 7.5, 50 mM sodium chloride, 2 mM magnesium chloride, and 2 mM DTT. Proteins were diluted in buffer then immediately combined and incubated at 25°C for 15 minutes before RNA was added and incubated at 25°C for another 15 minutes. Samples were stored on ice prior to grid freezing.

Samples were prepared for structural analysis using UltraAuFoil R1.2/1.3 300 mesh grids (Quantifoil) and a Vitrobot Mark IV (ThermoFisher Scientific). Grids were freshly glow discharged. Immediately before applying sample to grids, 0.5 μL n-Dodecyl-β-D-maltoside (DDM) was added to samples at a final concentration of 60 μM. 3.5 μL of sample was spotted onto grids before double-sided blotting and plunge freezing into liquid ethane.

### Cryo-EM Data Collection, Processing, and Model Building

SerialEM was used for data collection on a Titan Krios 300 keV transmission electron microscope (Thermo Fisher Scientific) [41]. Movies were collected on a K3 direct electron detector (Gatan) in CDS mode with a GIF quantum energy filter slit width of 20 eV and a stage tilt of 25°.

Patch motion correction, patch CTF estimation, particle picking, and particle extraction were performed using cryoSPARC v3.3.1 [42]. Image stacks were subjected to 2D classification, ab-initio model generation, 3D classification, and non-uniform refinement. The 3D reconstruction was used for variability analysis [31] and further 3D classification before a final non-uniform refinement and 3D reconstruction **(Table S1, Fig. S8 and S4).**

To build the PEDV polymerase complex coordinate model, a SARS-CoV-2 polymerase complex structure (PDB ID: 7CYQ) was docked into the cryo-EM map in Coot and regions of the protein (including whole chains of nsp8, nsp9, and nsp13) and RNA that lacked map density were removed [20, 43]. Mutation of protein chains to PEDV sequences, further model building, and validation was performed in Coot. The PEDV model was refined using real space refinement in Phenix [44]. Refinement in Phenix and model building/validation in Coot was an iterative process to produce a coordinate model. Final model adjustments were made with ISOLDE [45].

## Limitations of Study

We acknowledge that our study uses short RNA substrates, and short extension reactions and so does not model processive CoV RNA synthesis. This limits our ability to make conclusions on nsp8_2_’s importance for processive RNA synthesis. We believe studies using longer RNA substrates will allow our hypotheses and conclusions developed here to be evaluated in the context of processive replication but are beyond the scope of the current work.

Our work only studies the core polymerase complex (nsp7, nsp8, and nsp12) but our conclusions include hypotheses about interactions with other viral proteins (i.e., nsp13 and nsp14). While our current work is unable to directly assess these hypotheses, this work provides a foundation for further study.

## Supporting information

SuppMovie1

## Data Availability

The PEDV core polymerase complex electron density map has been deposited in the Electron Microscopy Data Bank (EMDB: 29779). The complex model has been deposited in the Protein Data Bank (PDB: 8G6R).

## Acknowledgements

We would like to thank the Cryo-EM Research Center in the Biochemistry Department at University of Wisconsin-Madison for technical and staff support. This work was supported by NIH/NIAID AI123498 and AI158463 and USDA WIS03099 grants to R.N.K, as well as NIH R35GM118110 grant to J.J.C. We would like to thank Dr. Olve Peersen, for his suggestions and insight for use of the nsp8-nsp7 fusion construct.

