## SupplementalData for "An alphacoronavirus polymerase structure reveals conserved co-factor functions"

| | |
|---|---|
| EMDB | 29779 |
| PDB | 8G6R |
| Microscope | Talos Arctica |
| Voltage (kV) | 200 |
| Detector | K3 direct electron detector (Gatan) |
| Dose Rate ($e^-$/pixel/sec) | 12 |
| Exposure Time (sec) | 3.5 |
| Electron Exposure ($e^-$/Å$^2$) | 60 |
| Frames (no.) | 60 |
| Defocus Values | -0.7 to -2.5 |
| Sample Tilt (°) | 25 |
| Data Collection Mode | EFTEM, Counting, CDS |
| Nominal Magnification | 105,000x |
| Pixel Size (Å) | 0.834 |
| Symmetry Imposed | C1 |
| Movies Collected (no.) | 1,261 |
| Initial Particle Images (no.) | 710,143 |
| Final Particle Images (no.) | 74,367 |
| Map Resolution (Å) – GSFSC | 3.3 |
| Initial Model Used (PDB ID) | 7CYQ |
| Non-hydrogen Atoms | 9,410 |
| Protein Residues | 1075 |
| Nuceic Acid Residues | 40 |
| R.M.S. Deviations | |
| Bond Lengths (Å) | 0.004 |
| Bond angles (°) | 0.556 |
| MolProbity Score | 1.64 |
| Clashscore | 7.19 |
| Ramachandran Plot | |
| Favored (%) | 96.34 |
| Allowed (%) | 3.66 |
| Disallowed (%) | 0 |

**Table S1: cryo-EM data collection and refinement.** Information provided is for the cryo-EM data collection, and processing that produced the electron density map of the PEDV polymerase complex. Additionally, validation statistics for the polymerase complex coordinate model built into the reconstruction are provided.

| | Desolvation Parameters | |
|---|---|---|
| **Sample** | **Capillary Temperature (°C)** | **In-source Trapping (V)** |
| PEDV nsp12+8+7 | 250 | -150 |
| PEDV A382R-nsp12+8+7 | 200 | -300 |
| PEDV V848R-nsp12+8+7 | 200 | -200 |
| PEDV nsp12+8+8L7 | 250 | -300 |
| SARS-CoV-2 nsp12+8+7 | 200 | -150 |
| SARS-CoV-2 L387R-nsp12+8+7 | 200 | -150 |
| SARS-CoV-2 T853R-nsp12+8+7 | 200 | -160 |
| SARS-CoV-2 nsp12+8+8L7 | 250 | -150 |

**Table S2, native mass spectrometry desolvation parameters:** Parameters listed were optimized for each sample to remove as many adducts as possible without disrupting the complex(es) of interest.

**Supplemental video 1, 3D variability analysis of PEDV core complex:** The movie displays a series of 3D volumes (in gray) determined by 3D variability analysis (cryoSPARC v3.3.1). The movie begins by oscillating between the extremes of variability within the volume series. Docking our PEDV core complex model (nsp12 – light purple, nsp7 – green, nsp8 – red, primer – neon green, template – dark purple) into the densities reveals the flexibility of dsRNA leaving the active site. We predict that this movement results in the lack of complete reconstruction of our dsRNA and $nsp8_1$ N-terminal extension in our final map and model.

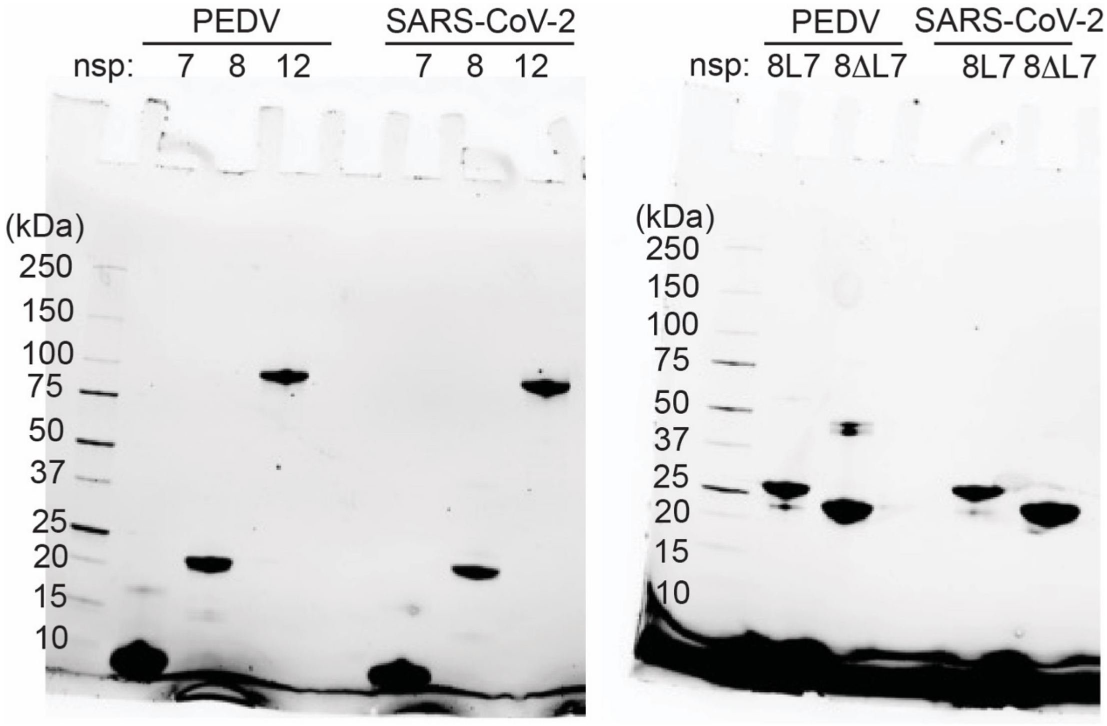


**Figure S1, SDS-PAGE of viral proteins:** Samples of purified proteins used for *in vitro* studies were run on pre-cast, stain-free, 4-20% SDS-PAGE gels (BioRad) and imaged using UV fluorescence. Expected molecular weights for different proteins are nsp7 – 9 kDa, nsp8 – 22 kDa, nsp12 – 110 kDa, nsp8L7 – 31 kDa, and nsp8ΔL7 – 25 kDa. The far-left lane of each gel is a protein ladder protein MWs labeled.

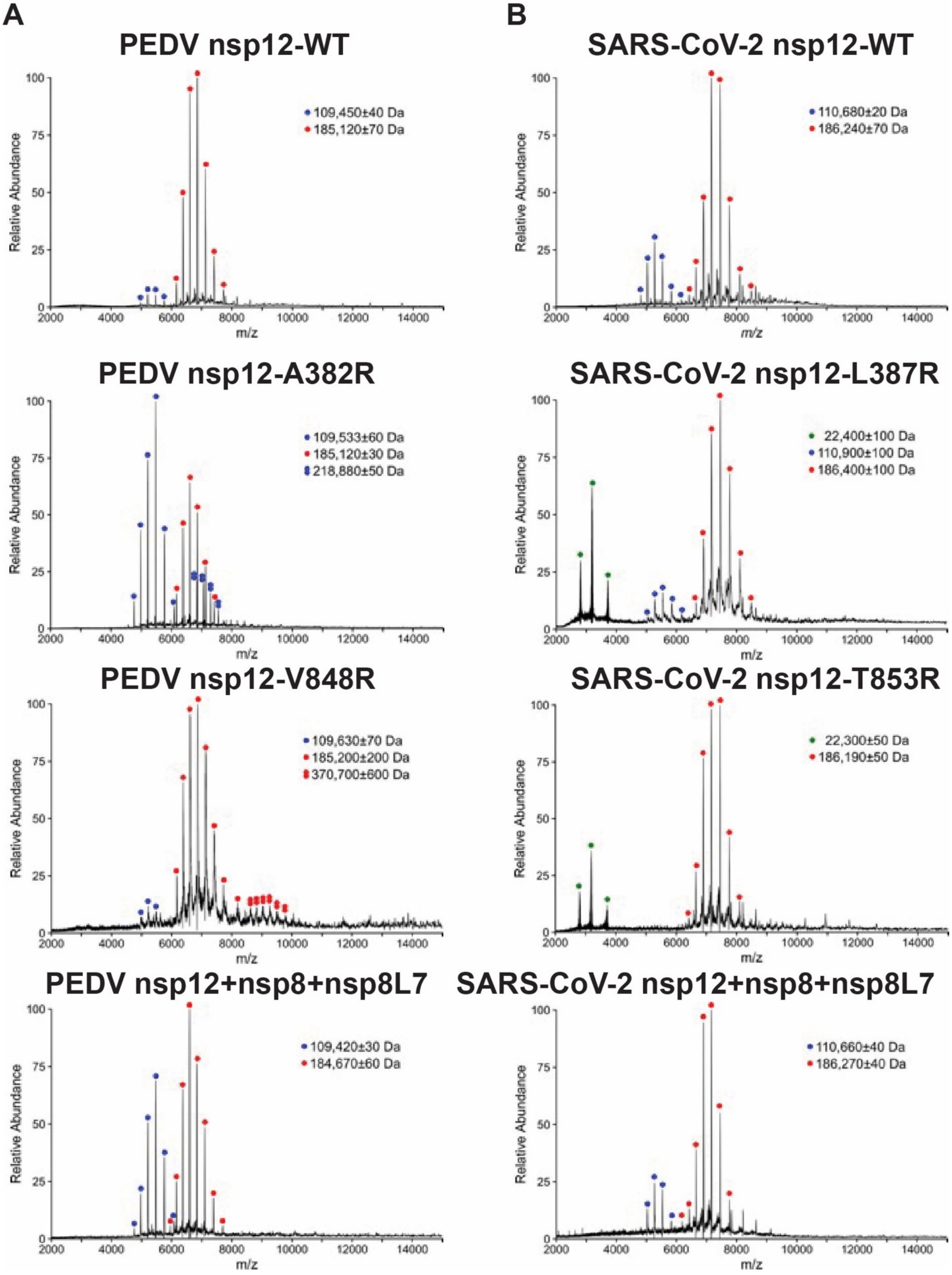


**Figure S2, native mass spectrometry of coronavirus polymerase complexes.** Native mass spectra for all coronavirus polymerase complexes tested with major mass populations labelled for each. Single and double red dots are monomeric and dimeric full intact complexes, respectively. Single and double blue dots are monomeric and dimeric solo nsp12, respectively. Green dots are free nsp8. **A)** SARS-CoV-2 complexes, top to bottom: wildtype, L387R, T853R, nsp8L7. **B)** PEDV complexes, top to bottom: wildtype, A382R, V848R, nsp8L7.

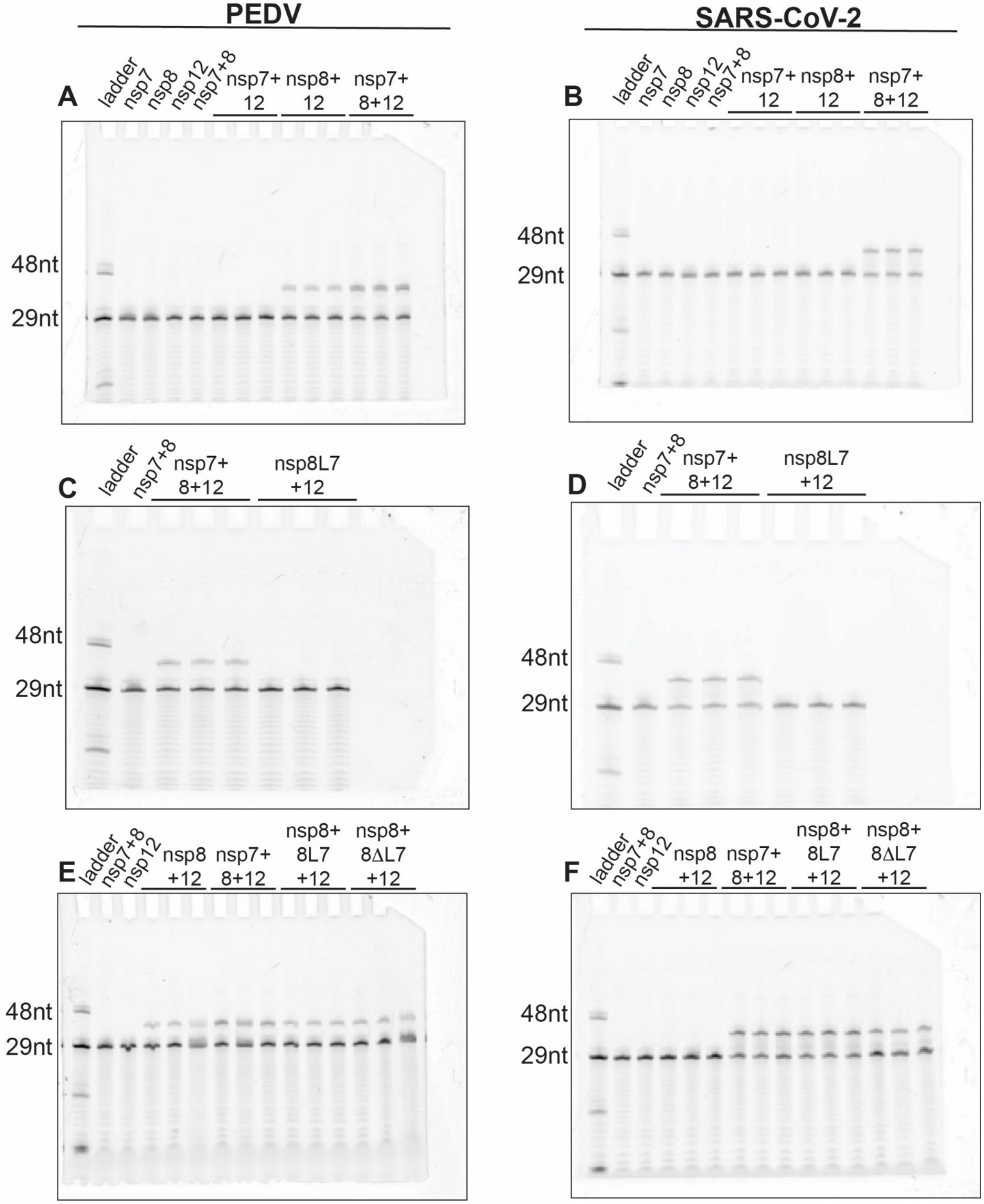


**Figure S3, full gel images for piecewise and fusion protein primer extensions:** Primer extensions assess the activity of nsp12 by its ability to extend an RNA primer (29 nucleotides or nt) to the length of its RNA template (38 nt) pair. **A)** PEDV piecewise primer extension revealing optimal nsp12 activity in the presence of both nsp7 and nsp8, and reduced activity with just nsp8. **B)** SARS2 piecewise primer extensions showing nsp12 is only active in the presence of both cofactors. Nsp8L7 does not individually activate nsp12 (**C** is PEDV, **D** is SARS-CoV-2). Nsp8L7 and nsp8ΔL7 activate nsp12 with free nsp8 present (**E** is PEDV, **F** is SARS-CoV-2), indicating polymerase activities requirement for $nsp8_1$, while $nsp8_2$'s helical extension is not essential for activity.

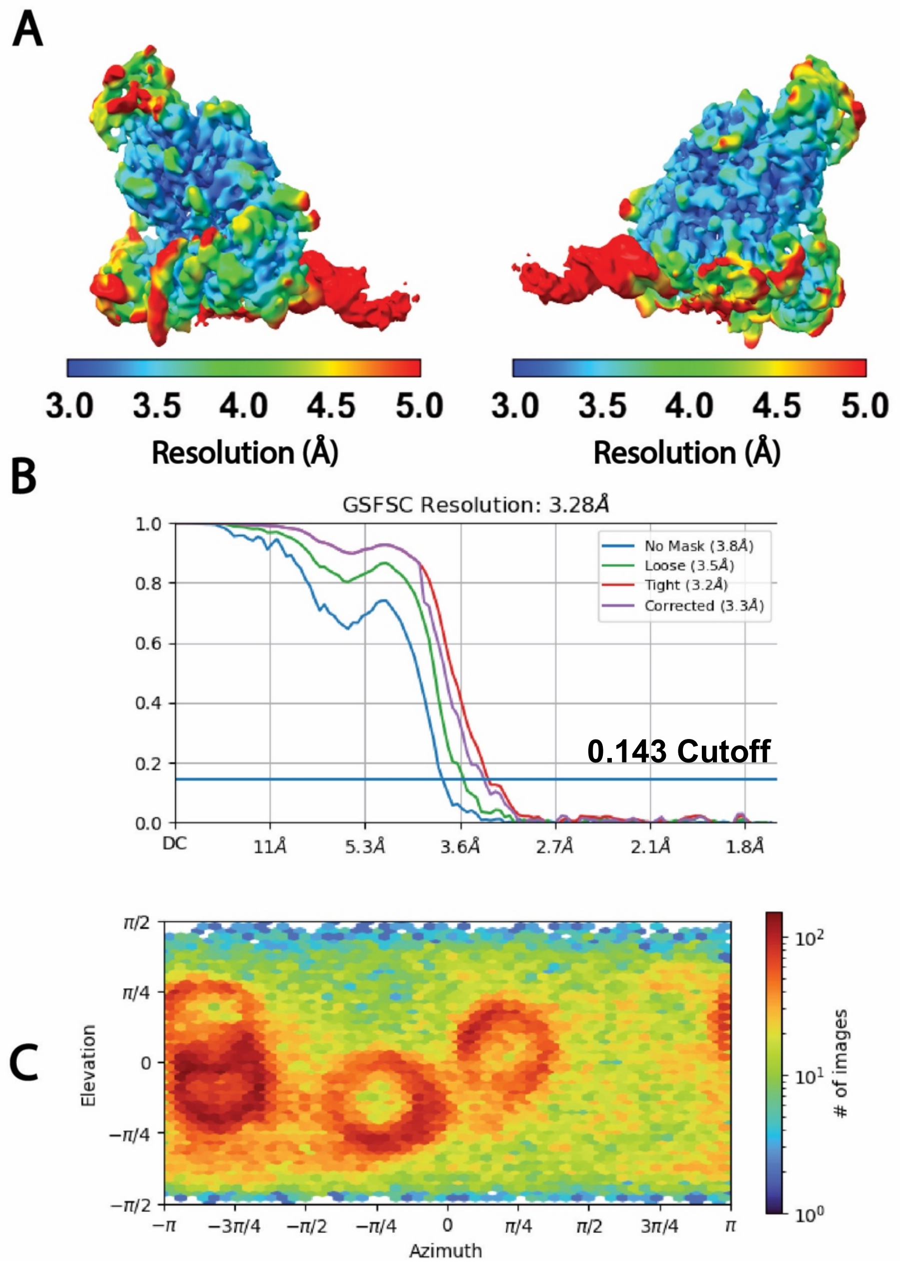


**Figure S4, cryo-EM validation:** A) Cryo-EM reconstruction filtered by local resolution according to the color key. B) Gold-standard Fourier shell correlation plot determined in cryoSPARC. The solid blue line marks the 0.143 cutoff. C) Particle orientation distribution plot for the final reconstruction. “# of images” indicates number of projections at a particular orientation (Elevation x Azimuth).

```
δ  HKU19        -LGYSSNQNNSYLNRVKGSS-DARLEPCTSDNRPDVVVRAFNIYNN--ATAGIFKSTKNN   56
   HKU11        --------NSPYLNRVTGSS-GARLEPQQPGVTPDAVKRAFHVHNN--TTSGIFLSTKTN   49
   PDCV         --------NSAYLNRVTGSS-DARLEPLQPGTQPDAVKRAFHVHND--TTSGIFLSTKSN   49
α  FCoV_65F     ----GTTVDQSYLNRVRGSS-AARLEPCN-GTDPDHVSRAFDIYNK--DVACIGKFLKTN   52
   HCoV_229E    ------SFDSSYLNRVRGSS-AARLEPCN-GTDIDYCVRAFDVYNK--DASFIGKNLKSN   50
   HCoV_NL63    ------SVDISYLNRARGSS-AARLEPCN-GTDIDKCVRAFDIYNK--NVSFLGKCLKMN   50
   HKU8         ------SLDNNYLNRVRGSS-AARLEPCN-GTEPEHVIRAFDIYNK--DVACIGKFVKVN   50
   PEDV         ------STDMAYLNRVRGSS-AARLEXCN-GTDTQHVYRAFDIYNK--DVACLGKFLKVN   50
γ  HKU22        ------SVPQNYLNRVRGLS-EARLRPCASGLLPDVVKRAFDLYNS--NTAGMYASLKHN   51
   IBV          SVAVASGFDKNYLNRVRGSS-EARLIPLANGCDPDVVKRAFDVCNK--ESAGMFQNLKRN   57
β  HCoV_OC43    ------SKDTNFLNRVRGASVDARLVPCASGLSTDVQLRAFDIYNA--SVAGIGLHLKVN   52
   HCoV_HKU1    ------SKDLNFLNRVRGTSVNARLVPCASGLSTDVQLRAFDICNT--NRAGIGLYYKVN   52
   MHV          ------SKDTNFLNRVRGTSVNARLVPCASGLDTDVQLRAFDICNA--NRAGIGLYYKVN   52
   MERS         ------SKDSNFLNRVRGSIVNARIEPCSSGLSTDVVFRAFDICNYKAKVAGIGKYYKTN   54
   HKU4         ------SKDTNFLNRVRGSSVNARLEPCSSGLTTDVVYRAFDICNFKARVAGIGKYYKTN   54
   HKU5         ------SKDSNFLNRVRGSIVNARIEPCASGLTTDVVFRAFDICNYKAKVAGIGKYYKTN   54
   HKU9         ------AKDECFLNRVRGTSGVARLVPLGSGVQPDIVLRAFDICNT--KVAGFGLHLKNN   52
   SARS-CoV-2   -----SADAQSFLNRVCGVS-AARLTPCGTGTSTDVVYRAFDIYND--KVAGFAKFLKTN   52
   SARS-CoV     -----SADASTFLNRVCGVS-AARLTPCGTGTSTDVVYRAFDIYNE--KVAGFAKFLKTN   52
                           :***. *    **:     .   :   ***.: *     : :    * *

δ  HKU19        CTRFKSTRPGSIINKPVRTIETFFVTKQCTENVFRAEEQCYNMLPKSIVSTDDKFSCVAY   116
   HKU11        CSRFRTTKQNLPLP-NKGSVELYFVSKQCSQQVFEIEETCYNMFDDSLKSTPEKFGVLAR   108
   PDCV         CARFKTTRSALPLP-NKGEVELYFVTKQCAAKVFEIEEECYNALSTELYTTDDTFGVLAK   108
α  FCoV_65F     CSRFRNLD----------KRDAYYVVKRCTKSVMDHEQVCYNDLKD--------SGAVAE   94
   HCoV_229E    CVRFKNVD----------KDDAFYIVKRCIKSVMDHEQSMYNLLKG--------CNAVAK   92
   HCoV_NL63    CVRFKNAD----------LKDGYFVIKRCTKSVMEHEQSMYNLLNF--------SGALAE   92
   HKU8         CVRFKNAD----------KHDAFYVVKRCTKSVMEHEQSIYDALKD--------CGAVSP   92
   PEDV         CVRLKNLD----------KHDAFYVVKRCTKSAMEHEQSIYSRLEK--------CGAVAE   92
γ  HKU22        CARFQELDENDD------EIDSFFVVKQTTPHNFEHEEKCYLDLKA---------DCVAV   96
   IBV          CARFQEVRDTED--GNLEYCDSYFVVKQTTPSNYEHEKACYEDLKS---------EVTAD   106
β  HCoV_OC43    CCRFQRVDENGD------KLDQFFVVKRTDLTIYNREMKCYERVKD--------CKFVAE   98
   HCoV_HKU1    CCRFQRIDDDGN------KLDKFFVVKRTNLEVYNKEKTYYELTKS--------CGVVAE   98
   MHV          CCRFQRVDEDGN------KLDKFFVVKRTNLEVYNKEKECYELTKE--------CGVVAE   98
   MERS         TCRFVELDDQGH------HLDSYFVVKRHTMENYELEKHCYDLLRD--------CDAVAP   100
   HKU4         TCRFVQVDDEGH------KLDSYFIVKRHTMSNYELEKRCYDLLKD--------CDAVAI   100
   HKU5         TCRFVEVDDEGH------RLDSFFVVKRHTMENYELEKRCYDLVKD--------CDAVAV   100
   HKU9         CCRYQELDADGN------QLDSYFVVKRHTESNYLLEQRCYEKLKD--------CDVVAR   98
   SARS-CoV-2   CCRFQEKDEDDN------LIDSYFVVKRHTFSNYQHEETIYNLLKD--------CPAVAK   98
   SARS-CoV     CCRFQEKDEEGN------LLDSYFVVKRHTMSNYQHEETIYNLVKD--------CPAVAV   98
                  *                 : ::: *:        *   *                :

δ  HKU19        HDFFKFDG----VNNVVRRHLTKYTLLDLVYALRHLSTS-QEIIQEILTTMCGTSEE---   168
   HKU11        TEFFKFDK----IPNVNRQFLTKYTLLDLAYALRHLSTS-RDVIKEILITICGTTEE---   160
   PDCV         TEFFKFDK----IPNVNRQYLTKYTLLDLAYALRHLSTS-KDVIQEILITMCGTPED---   160
α  FCoV_65F     HDFFLYKEGRCEFGNVARKDLTKYTMMDLCYAIRNFDEKNCEVLKEILVTLGACNES---   151
   HCoV_229E    HDFFTWHEGRTIYGNVSRQDLTKYTMMDLCFALRNFDEKDCGVFKEILVLTGCCNTD---   149
   HCoV_NL63    HDFFTWKDGRVIYGNVSRHNLTKYTMMDLVYAMRNFDEQNCDVLKEVLVLTGCCDNS---   149
   HKU8         HDFFVWKDGRSVYGNIARHDLTKYTMMDLVHALRNFDEKNCETLKEILVISGACDSS---   149
   PEDV         HDFFTWKDGRAIYGNVCRKDLTEYTMMDLCYALRNFDENNCDVLKSILIKVGACEES---   149
γ  HKU22        HDFFRFE-G---MYNICRQRLTKYTMMDLCYAFRHFDPNDCDVLKEILVVKGCCEWD---   149
   IBV          HDFFVFNKN---IYNISRQRLTKYTMMDFCYALRHFDPKDCEVLKEILVTYGCIEDYHPK   163
β  HCoV_OC43    HDFFTFDVEGSRVPHIVRKDLTKYTMLDLCYALRHFDRNDCMLLCDILSIYAGCEQS---   155
   HCoV_HKU1    HDFFTFDIDGSRVPHIVRRNLSKYTMLDLCYALRHFDRNDCSILCEILCEYADCKES---   155
   MHV          HEFFTFDVEGSRVPHIVRKDLSKFTMLDLCYALRHFDRNDCSTLKEILLTYAECEES---   155
   MERS         HDFFIFDVDKVKTPHIVRQRLTEYTMMDLVYALRHFDQN-SEVLKAILVKYGCCDVT---   156
   HKU4         HDFFIFDVDKTKTPHIVRQSLTEYTMMDLVYALRHFDQNNCEVLKSILVKYGCCEQS---   157
   HKU5         HDFFIFDVDKVKTPHIVRQRLTEYTMMDLVYALRHFDQNNCEVLKSILVKYGCCDAS---   157
   HKU9         HDFFKFNIEGVMTPHVSRERLTKYTMADLVYSLRHFDNNNCDTLKEILVLRGCCTVD---   155
   SARS-CoV-2   HDFFKFRIDGDMVPHISRQRLTKYTMADLVYALRHFDEGNCDTLKEILVTYNCCDDD---   155
   SARS-CoV     HDFFKFRVDGDMVPHISRQRLTKYTMADLVYALRHFDEGNCDTLKEILVTYNCCDDD---   155
                 :** :          :: *. *:::*: *: .::*::.      :  :*
```

NiRAN
Motif $A_N$

NiRAN
Motif $B_N$

```
δ  HKU19        WF--VDGWYDPIENPTFYDEFHKLGSLINNCVVMANKFADTCKTVGLVGILTADNQDLGG   226
δ  HKU11        WF--GDSWFDPIENPTFYREFHKLGSVLNRCVLNANAFAKACSELGIVGILTPDNQDLLG   218
δ  PDCV         WF--GENWFDPIENPSFYKEFHKLGDILNRCVLNANKFASACIDAGLVGILTPDNQDLLG   218
α  FCoV_65F     -FFENKDWFDPVENEAIHEVYARLGPIVANAMLKCVAFCDAIVEKGYIGIITLDNQDLNG   210
α  HCoV_229E    -YFEMKNWFDPIENEDIHRVYAALGKVVANAMLKCVAFCDEMVLKGVVGVLTLDNQDLNG   208
α  HCoV_NL63    -YFDSKGWYDPVENEDIHRVYASLGKIVARAMLKCVALCDAMVAKGVVGVLTLDNQDLNG   208
α  HKU8         -YFDNKNWYDPVENEDIHRVYAKLGCVVANAMLKCVALCDAMVAKGVVGVLTLDNQDLNG   208
α  PEDV         -YFNNKVWFDPVENEDIHRVYALLGTIVSRAMLKCVKFCDAMVEQGIVGVVTLDNQDLNG   208
γ  HKU22        -YFDQPNWYDPVENPDWFSLISRLGPIFQRALIKVAEFCDLMVEKGYIGVVTLDNQDLNG   208
γ  IBV          WFEENKDWYDPIENPKYYAMLAKMGPIVRRALLNAIEFGNLMVEKGYVGVITLDNQDLNG   223
β  HCoV_OC43    -YFTKKDWYDFVENPDIINVYKKLGPIFNRALVSATEFADKLVEVGLVGVLTLDNQDLNG   214
β  HCoV_HKU1    -YFSKKDWYDFVENPDIINIYKKLGPIFNRALLNTVIFADTLVEVGLVGVLTLDNQDLYG   214
β  MHV          -YFQKKDWYDFVENPDIINVYKKLGPIFNRALLNTAKFADALVEAGLVGVLTLDNQDLYG   214
β  MERS         -YFENKLWFDFVENPSVIGVYHKLGERVRQAILNTVKFCDHMVKAGLVGVLTLDNQDLNG   215
β  HKU4         -YFDNKLWFDFVENPSVIGVYHKLGERIRQAMLNTVKMCDHMVKSGLVGVLTLDNQDLNG   216
β  HKU5         -YFDNKLWFDFVENPNVISVYHKLGERIRQAVLNTVKFCDQMVKSGLVGVLTLDNQDLNG   216
β  HKU9         -YFDRKDWYDPVENPDIIRVYHKLGETVRKAVLSAVKMADAMVEQGLIGVLTLDNQDLNG   214
β  SARS-CoV-2   -YFNKKDWYDFVENPDILRVYANLGERVRQALLKTVQFCDAMRNAGIVGVLTLDNQDLNG   214
β  SARS-CoV     -YFNKKDWYDFVENPDILRVYANLGERVRQSLLKTVQFCDAMRDAGIVGVLTLDNQDLNG   214
                 :     *:* :**          :*  . ..::    : .     * :*::* ***** *

δ  HKU19        QIYDFGDFVVTQPGNGCIEMDAYLSYIMPSMSMTHMLKCECLDD----NGSYKDYSIYQY   282
δ  HKU11        QIYDFGDFIITQPGNGCVDLSSYYSYLMPIMSMTHMLKCECYDN----DGNEIDYDGFQY   274
δ  PDCV         QIYDFGDFIITQPGNGCVDLASYYSYLMPIMSMTHMLKCECMDS----DGNPLEYDGFQY   274
α  FCoV_65F     NFYDFGDFVKTAPGFGCACVTSYYSYMMPLMGMTSCLESENFVKSDIYGSDYKQYDLLAY   270
α  HCoV_229E    NFYDFGDFVLCPPGMGIPYCTSYYAYMMPVMGMTNCLASECFMKSDIFGQDFKTFDLLKY   268
α  HCoV_NL63    NFYDFGDFVVSLPNMGVPCCTSYYSYMMPIMGLTNCLASECFVKSDIFGSDFKTFDLLKY   268
α  HKU8         NFYDFGDFTIGIPGVGVPLATSYYSYLMPVMGMTNCLARECFVKSEIFGSDFKTYDLLEY   268
α  PEDV         DFYDFGDFTCSIKGMGIPICTSYYSYMMPVMGMTNCLASECFVKSDIFGEDFKSYDLLEY   268
γ  HKU22        NFYDFGDFKKVLPGCGVPVTTSYYSYMMPCLTACDALASERFFEFKA-TSGYKQYDLTKY   267
γ  IBV          KFYDFGDFQKTAPGAGVPVFDTYYSYMMPIIAMTDALAPERYFEYDV-HKGYKSYDLLKY   282
β  HCoV_OC43    KWYDFGDYVIAAPGCGVAIADSYYSYIMPMLTMCHALDCELYV-----NNAYRLFDLVQY   269
β  HCoV_HKU1    QWYDFGDFIQTAPGFGVAVADSYYSYMMPMLTMCHVLDCELFV-----NDSYRQFDLVQY   269
β  MHV          QWYDFGDFVKTVPGCGVAVADSYYSYMMPMLTMCHALDSELFV-----NGTYREFDLVQY   269
β  MERS         KWYDFGDFVITQPGSGVAIVDSYYSYLMPVLSMTDCLAAETHRDCDF-NKPLIEWPLTEY   274
β  HKU4         KWYDFGDFVITQPGAGVAIVDSYYSYLMPVLSMTNCLAAETHKDCDF-NKPLIEWPLLEY   275
β  HKU5         KWYDFGDFVITQPGAGVAIVDSYYSYLMPVLSMTNCLAAETHRDCDL-TKPLIEWPLLEY   275
β  HKU9         QWYDFGDFIEGPAGAGVAVMDTYYSLAMPIYTMTNILAAECHVSGDL-CNLKRVLDIFKY   273
β  SARS-CoV-2   NWYDFGDFIQTTPGSGVPVVDSYYSLLMPILTLTRALTAESHVDTDL-TKPYIKWDLLKY   273
β  SARS-CoV     NWYDFGDFVQVAPGCGVPIVDSYYSLLMPILTLTRALAAESHMDADL-AKPLIKWDLLKY   273
                . *****:     . *     :* :  **        *  *                    *

δ  HKU19        DFTDYKMELFNKYFRHWSQTYHPNCVDCVDDRCIVHCANFNILFAMCLPNTCFGNLCSQA   342
δ  HKU11        DFTDFKLSLFSKYFTYWDRPYHPNTVDCPDDRCVLHCANFNVLFAMCIPSTAFGNLCSQA   334
δ  PDCV         DFTDFKLGLFEKYFKYWDRPYHPNTVECPDDRCVLHCANFNVLFAMCIPNTAFGNLCSRA   334
α  FCoV_65F     DFTDHKEKLFEKYFKHWDRTYHPNCSDCTSDDCIIHCANFNTLFSMTIPNTAFGPLVRKV   330
α  HCoV_229E    DFTEHKEVLFNKYFKYWGQDYHPDCVDCHDEMCILHCSNFNTLFATTIPNTAFGPLCRKV   328
α  HCoV_NL63    DFTEHKENLFNKYFKHWSFDYHPNCSDCYDDMCVIHCANFNTLFATTIPGTAFGPLCRKV   328
α  HKU8         DFTEHKLGLFNKYFKHWDLDYHPNCSDCYDEMCVIHCANFNALFATTIPDTSFGPLCRKV   328
α  PEDV         DFTEHKTALFNKYFKYWGLQYHPNCVDCSDEQCIVHCANFNTLFSTTIPITAFGPLCRKC   328
γ  HKU22        DFTEEKLQLFMKYFKYWDRTYHPNCVECIDDRCLIHCANFNILFATLFPQTAFGCLCKRV   327
γ  IBV          DYTEEKQDLFQKYFKYWDQEYHPNCRDCSDDRCLIHCANFNILFSTLVPQTSFGNLCRKV   342
β  HCoV_OC43    DFTDYKLELFNKYFKHWSMPYHPNTVDCQDDRCIIHCANFNILFSMVLPNTCFGPLVRQI   329
β  HCoV_HKU1    DFTDYKLELFNKYFKYWGMKYHPNTVDCDNDRCIIHCANFNILFSMVLPNTCFGPLVRQI   329
β  MHV          DFTDFKLELFTKYFKHWSMTYHPNTCECEDDRCIIHCANFNILFSMVLPKTCFGPLVRQI   329
β  MERS         DFTDYKVQLFEKYFKYWDQTYHANCVNCTDDRCVLHCANFNVLFAMTMPKTCFGPIVRKI   334
β  HKU4         DYTDYKIGLFNKYFKYWDQTYHPNCVNCSDDRCILHCANFNVLFSMVLPNTSFGPIVRKI   335
β  HKU5         DYTDYKIGLFEKYFKXWDQQYHPNCVNCTDDRCVLHCANFNVLFSMTLPGTSFGPIVRKI   335
β  HKU9         YYTQFKYSLFSNYFKYWDMQYHPNCVACADDRCILHCANFNILFSMVLPNTSFGPLVQKI   333
β  SARS-CoV-2   DFTEERLKLFDRYFKYWDQTYHPNCVNCLDDRCILHCANFNVLFSTVFPPTSFGPLVRKI   333
β  SARS-CoV     DFTEERLCLFDRYFKYWDQTYHPNCINCLDDRCILHCANFNVLFSTVFPPTSFGPLVRKI   333
                  :*: :  ** .**  *.  ** :   * .: *::**:*** **:  .* *.** :  :
```

NiRAN
Motif $C_N$

NiRAN
Motif $C_N$

αCoV unique loop,
PEDV 249-268

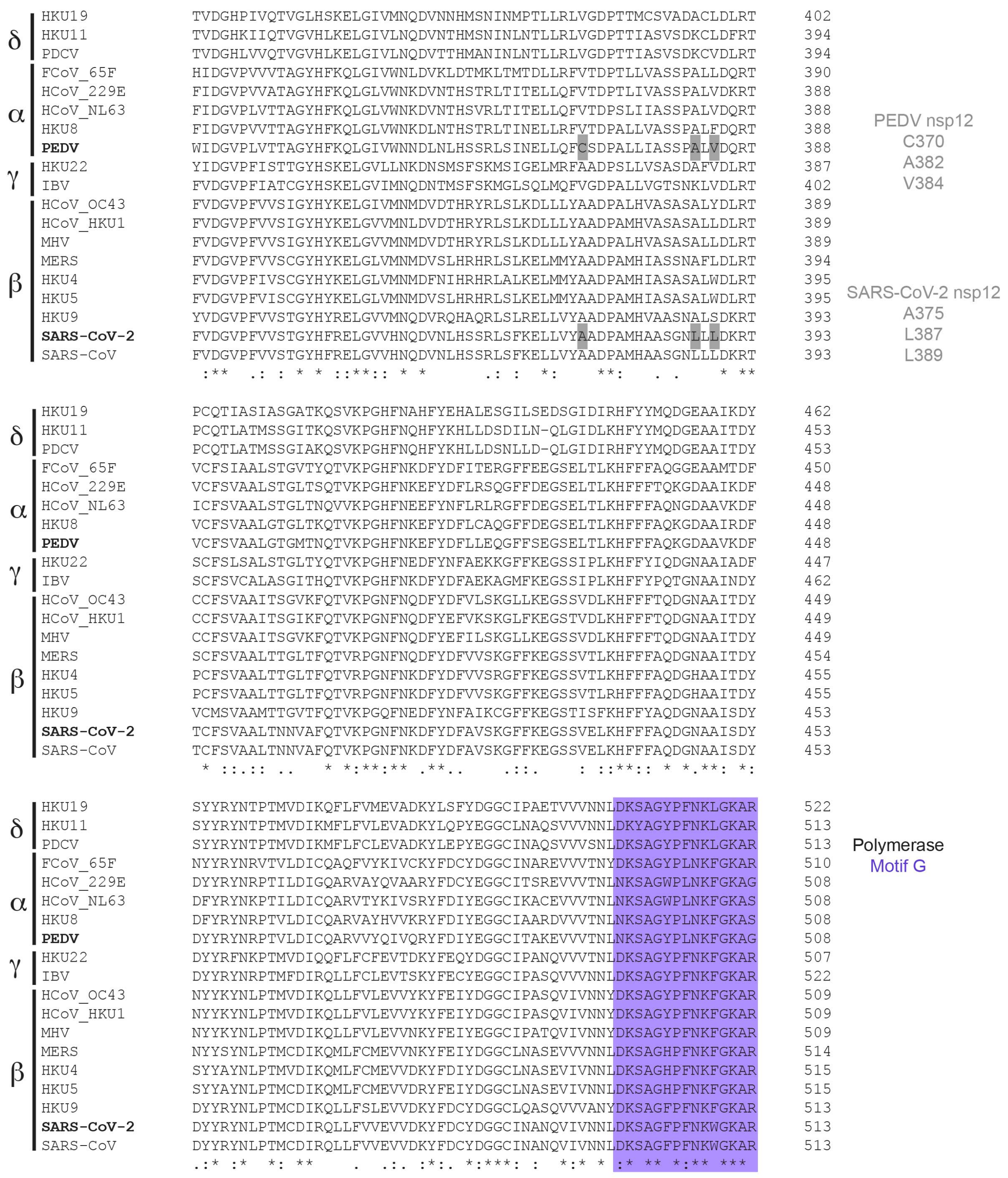

δ
α
γ
β
HKU19
HKU11
PDCV
FCoV_65F
HCoV_229E
HCoV_NL63
HKU8
PEDV
HKU22
IBV
HCoV_OC43
HCoV_HKU1
MHV
MERS
HKU4
HKU5
HKU9
SARS-CoV-2
SARS-CoV
PEDV nsp12
C370
A382
V384
SARS-CoV-2 nsp12
A375
L387
L389
Polymerase
Motif G

```
δ  HKU19        NYYD-LTYAEQNAMFEYTKRNVLPTVTQMNLKYAISAKDRARTVAGVSIISTMTNRQYHQ   581
δ  HKU11        NYYD-MTYAEQNQLFEYTKRNVLPTLTQMNLKYAISAKDRARTVAGVSIISTMTNRQYHQ   572
δ  PDCV         NYYD-MTHAEQNQLFEYTKRNVLPTLTQMNLKYAISAKDRARTVAGVSIISTMTNRQYHQ   572
α  FCoV_65F     LYYETLSYEEQDAIFALTKRNVLPTMTQMNLKYAISGKARARTVGGVSLLSTMTTRQYHQ   570
α  HCoV_229E    LYYESISYEEQDAIFLLTKRNILPTMTQLNLKYAISGKERARTVGGVSLLATMTTRQFHQ   568
α  HCoV_NL63    LYYESISYEEQDALFALTKRNVLPTMTQLNLKYAISGKERARTVGGVSLLSTMTTRQYHQ   568
α  HKU8         LYYESLSYEEQDALYALTKRNVLPTMTQLNLKYAISGKERARTVGGVSLLSTMTTRQFHQ   568
α  PEDV         LYYESLSYEEQDELYAYTKRNILPTMTQLNLKYAISGKERARTVGGVSLLSTMTTRQYHQ   568
γ  HKU22        LYYESLSYAEQDQLFELTKRNILPTITQINMKYAISAKSRARTVAGVSILSTMTNRQFHQ   567
γ  IBV          LYYE-MSLEEQDQLFESTKKNVLPTITQMNLKYAISAKNRARTVAGVSILSTMTNRQFHQ   581
β  HCoV_OC43    LYYEALSFEEQDEIYAYTKRNVLPTLTQMNLKYAISAKNRARTVAGVSILSTMTGRMFHQ   569
β  HCoV_HKU1    LYYEALSFEEQNEIYAYTKRNVLPTLTQMNLKYAISAKNRARTVAGVSILSTMTGRMFHQ   569
β  MHV          LYYEALSFEEQDEIYAYTKRNVLPTLTQMNLKYAISAKNRARTVAGVSILSTMTGRMFHQ   569
β  MERS         VYYESMSYQEQDELFAMTKRNVIPTMTQMNLKYAISAKNRARTVAGVSILSTMTNRQYHQ   574
β  HKU4         VYYESMSYQEQDELFAVTKRNVLPTITQMNLKYAISAKNRARTVAGVSILSTMTNRQYHQ   575
β  HKU5         VYYESLSYQEQDELFAMTKRNVLPTITQMNLKYAISAKNRARTVAGVSILSTMTNRQYHQ   575
β  HKU9         LYYESLSYADQDELFAYTKRNVLPTITQMNLKYAISAKNRARTVAGVSIASTMTNRQFHQ   573
β  SARS-CoV-2   LYYDSMSYEDQDALFAYTKRNVIPTITQMNLKYAISAKNRARTVAGVSICSTMTNRQFHQ   573
β  SARS-CoV     LYYDSMSYEDQDALFAYTKRNVIPTITQMNLKYAISAKNRARTVAGVSICSTMTNRQFHQ   573
                 **: ::  :*: ::  **:*::**:**:*:*****.* *****.***: :*** * :**
```

Polymerase Motif F

```
δ  HKU19        KLLKSISVARNQTIVIGTTKFYGGWDNMLRNLMANINNPKLAGWDYPKCDRSMPNLLRIT   641
δ  HKU11        KMLKSISLARNQTIVIGTTKFYGGWDNMLRRLMNGINNPILVGWDYPKCDRSMPNMLRIA   632
δ  PDCV         KMLKSISLARNQTIVIGTTKFYGGWDNMLRRLMCNINNPILVGWDYPKCDRSMPNMLRIA   632
α  FCoV_65F     KHLKSIAATRNATVVIGTTKFYGGWDNMLKNLMRDVDNGCLMGWDYPKCDRALPNMIRMA   630
α  HCoV_229E    KCLKSIVATRNATVVIGTTKFYGGWDNMLKNLMADVDDPKLMGWDYPKCDRAMPSMIRML   628
α  HCoV_NL63    KHLKSIVNTRNATVVIGTTKFYGGWNNMLRTLIDGVENPMLMGWDYPKCDRALPNMIRMI   628
α  HKU8         KHLKSIVNTRNATVVIGTTKFYGGWDNMLRNLMDGVDNACLMGWDYPKCDRALPNMIRMI   628
α  PEDV         KHLKSIVNTRGASVVIGTTKFYGGWDNMLKNLIDGVENPCLMGWDYPKCDRALPNMIRMI   628
γ  HKU22        KCLKSIVNTRNATVVIGTTKFYGGWDNMLRNLMRGVEDPVLMGWDYPKCDRAMPSLLRLS   627
γ  IBV          KILKSIVNTRNAPVVIGTTKFYGGWDNMLRNLIQGVEDPILMGWDYPKCDRAMPNLLRIA   641
β  HCoV_OC43    KCLKSIAATRGVPVVIGTTKFYGGWDDMLRRLIKDVDNPVLMGWDYPKCDRAMPNLLRIV   629
β  HCoV_HKU1    KCLKSIAATRGVPVVIGTTKFYGGWDDMLRHLIKDVDNPVLMGWDYPKCDRAMPNILRIV   629
β  MHV          KCLKSIAATRGVPVVIGTTKFYGGWDDMLRRLIKDVDSPVLMGWDYPKCDRAMPNILRIV   629
β  MERS         KMLKSMAATRGATCVIGTTKFYGGWDFMLKTLYKDVDNPHLMGWDYPKCDRAMPNMCRIF   634
β  HKU4         KMLKSMAATRGATCVIGTTKFYGGWDFMLKTLYKDVESPHLMGWDYPKCDRAMPNMCRIL   635
β  HKU5         KMLKSMAATRGSTCVIGTTKFYGGWDFMLKTLYKDVDNPHLMGWDYPKCDRAMPNMCRIF   635
β  HKU9         KMLKSIAAARGASVVIGTTKFYGGWNRMLRTLCEGVDNPHLMGWDYPKCDRAMPNLLRIF   633
β  SARS-CoV-2   KLLKSIAATRGATVVIGTSKFYGGWHNMLKTVYSDVENPHLMGWDYPKCDRAMPNMLRIM   633
β  SARS-CoV     KLLKSIAATRGATVVIGTSKFYGGWHNMLKTVYSDVETPHLMGWDYPKCDRAMPNMLRIM   633
                * ***:  :*.   ****:******. **: :  .::  * *********::*.: *:
```

Polymerase Motif A

```
δ  HKU19        SSLLLARKH-ACCTHSQRFYRLANECAQVLSEIVVSGNVMYVKPGGTSSGDATTAYANSV   700
δ  HKU11        ASCLLARKH-TCCNQSQRFYRLANECCQVLSEVVVSGNNLYVKPGGTSSGDATTAYANSV   691
δ  PDCV         ASCLLARKH-TCCNQSQRFYRLANECCQVLSEVVVSGNNLYVKPGGTSSGDATTAYANSV   691
α  FCoV_65F     SAMVLGSKHIGCCTHSDRFYRLSNELAQVLTEVVHCTGGFYIKPGGTTSGDGTTAYANSA   690
α  HCoV_229E    SAMILGSKHVTCCTASDKFYRLSNELAQVLTEVVYSNGGFYFKPGGTTSGDATTAYANSV   688
α  HCoV_NL63    SAMVLGSKHVNCCTATDRFYRLGNELAQVLTEVVYSNGGFYFKPGGTTSGDASTAYANSI   688
α  HKU8         SAMILGSKHVNCCTNSDRYYRLCNELAQVLTEVVYSNGGFYMKPGGTTSGDATTAYANSV   688
α  PEDV         SAMILGSKHTTCCSSTDRFFRLCNELAQVLTEVVYSNGGFYLKPGGTTSGDATTAYANSV   688
γ  HKU22        ASLILARRHKGCCDWNERIYRLANEAAQVLSEVALSNGGLYVKPGGTSSGDATTAYANSA   687
γ  IBV          ASLVLARKHTNCCTWSERVYRLYNECAQVLSETVLATGGIYVKPGGTSSGDATTAYANSV   701
β  HCoV_OC43    SSLVLARKHETCCSQSDRFYRLANECAQVLSEIVMCGGCYYVKPGGTSSGDATTAFANSV   689
β  HCoV_HKU1    SSLVLARKHEFCCSHGDRFYRLANECAQVLSEIVMCGGCYYVKPGGTSSGDATTAFANSV   689
β  MHV          SSLVLARKHDSCCSHTDRFYRLANECAQVLSEIVMCGGCYYVKPGGTSSGDATTAFANSV   689
β  MERS         ASLILARKHGTCCTTRDRFYRLANECAQVLSEYVLCGGGYYVKPGGTSSGDATTAYANSV   694
β  HKU4         ASLILARKHSTCCTNSDRFYRLANECAQVLSEYVLCGGGYYVKPGGTSSGDATTAYANSV   695
β  HKU5         ASLILARKHSTCCTNTDRFYRLANECAQVLSEYVLCGGGYYVKPGGTSSGDATTAYANSV   695
β  HKU9         ASLILARKHSTCCNASERFYRLANECAQVLSEMVLCGGGFYVKPGGTSSGDSTTAYANSV   693
β  SARS-CoV-2   ASLVLARKHTTCCSLSHRFYRLANECAQVLSEMVMCGGSLYVKPGGTSSGDATTAYANSV   693
β  SARS-CoV     ASLVLARKHNTCCNLSHRFYRLANECAQVLSEMVMCGGSLYVKPGGTSSGDATTAYANSV   693
                :: :*. :*  **   .: :** ** .***:* . . .  *.*****:***.:**:***
```

Polymerase Motif B

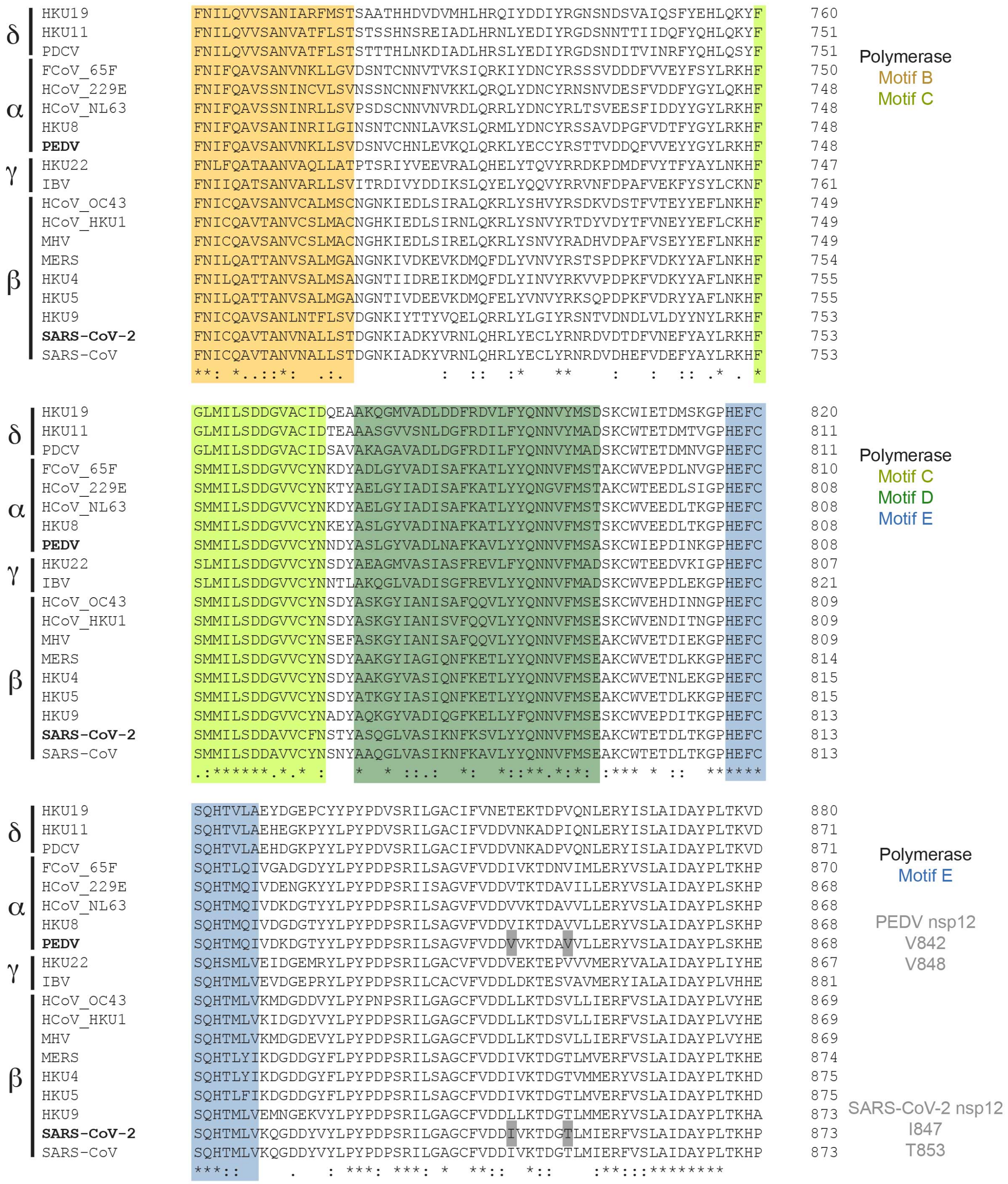
δ
α
γ
β
HKU19 FNILQVVSANIARFMSTSAATHHDVDVMHLHRQIYDDIYRGNSNDSVAIQSFYEHLQKYF 760
HKU11 FNILQVVSANVATFLSTSTSSHNSREIADLHRNLYEDIYRGDSNNTTIIDQFYQHLQKYF 751
PDCV FNILQVVSANVATFLSTSTTTHLNKDIADLHRSLYEDIYRGDSNDITVINRFYQHLQSYF 751
FCoV_65F FNIFQAVSANVNKLLGVDSNTCNNVTVKSIQRKIYDNCYRSSSVDDDFVVEYFSYLRKHF 750
HCoV_229E FNIFQAVSSNINCVLSVNSSNCNNFNVKKLQRQLYDNCYRNSNVDESFVDDFYGYLQKHF 748
HCoV_NL63 FNIFQAVSSNINRLLSVPSDSCNNVNVRDLQRRLYDNCYRLTSVEESFIDDYYGYLRKHF 748
HKU8 FNIFQAVSANINRILGINSNTCNNLAVKSLQRMLYDNCYRSSAVDPGFVDTFYGYLRKHF 748
PEDV FNIFQAVSANVNKLLSVDSNVCHNLEVKQLQRKLYECCYRSTTVDDQFVVEYYGYLRKHF 748
HKU22 FNLFQATAANVAQLLATPTSRIYVEEVRALQHELYTQVYRRDKPDMDFVYTFYAYLNKHF 747
IBV FNIIQATSANVARLLSVITRDIVYDDIKSLQYELYQQVYRRVNFDPAFVEKFYSYLCKNF 761
HCoV_OC43 FNICQAVSANVCALMSCNGNKIEDLSIRALQKRLYSHVYRSDKVDSTFVTEYYEFLNKHF 749
HCoV_HKU1 FNICQAVTANVCSLMACNGHKIEDLSIRNLQKRLYSNVYRTDYVDYTFVNEYYEFLCKHF 749
MHV FNICQAVSANVCSLMACNGHKIEDLSIRELQKRLYSNVYRADHVDPAFVSEYYEFLNKHF 749
MERS FNILQATTANVSALMGANGNKIVDKEVKDMQFDLYVNVYRSTSPDPKFVDKYYAFLNKHF 754
HKU4 FNILQATTANVSALMSANGNTIIDREIKDMQFDLYINVYRKVVPDPKFVDKYYAFLNKHF 755
HKU5 FNILQATTANVSALMGANGNTIVDEEVKDMQFELYVNVYRKSQPDPKFVDRYYAFLNKHF 755
HKU9 FNICQAVSANLNTFLSVDGNKIYTTYVQELQRRLYLGIYRSNTVDNDLVLDYYNYLRKHF 753
SARS-CoV-2 FNICQAVTANVNALLSTDGNKIADKYVRNLQHRLYECLYRNRDVDTDFVNEFYAYLRKHF 753
SARS-CoV FNICQAVTANVNALLSTDGNKIADKYVRNLQHRLYECLYRNRDVDHEFVDEFYAYLRKHF 753
**: *..::*:  .:.          :  ::  :*    **     :    :   :: .* . *
Polymerase
Motif B
Motif C
δ
α
γ
β
HKU19 GLMILSDDGVACIDQEAAKQGMVADLDDFRDVLFYQNNVYMSDSKCWIETDMSKGPHEFC 820
HKU11 GLMILSDDGVACIDTEAAASGVVSNLDGFRDILFYQNNVYMADSKCWTETDMTVGPHEFC 811
PDCV GLMILSDDGVACIDSAVAKAGAVADLDGFRDILFYQNNVYMADSKCWTETDMNVGPHEFC 811
FCoV_65F SMMILSDDGVVCYNKDYADLGYVADISAFKATLYYQNNVFMSTAKCWVEPDLNVGPHEFC 810
HCoV_229E SMMILSDDGVVCYNKTYAELGYIADISAFKATLYYQNGVFMSTAKCWTEEDLSIGPHEFC 808
HCoV_NL63 SMMILSDDGVVCYNKDYAELGYIADISAFKATLYYQNNVFMSTSKCWVEEDLTKGPHEFC 808
HKU8 SMMILSDDGVVCYNKEYASLGYVADINAFKATLYYQNNVFMSTSKCWVEEDLTKGPHEFC 808
PEDV SMMILSDDGVVCYNNDYASLGYVADLNAFKAVLYYQNNVFMSASKCWIEPDINKGPHEFC 808
HKU22 SLMILSDDGVVCYNSDYAEAGMVASIASFREVLFYQNNVFMADSKCWTEEDVKIGPHEFC 807
IBV SLMILSDDGVVCYNNTLAKQGLVADISGFREVLYYQNNVFMADSKCWVEPDLEKGPHEFC 821
HCoV_OC43 SMMILSDDGVVCYNSDYASKGYIANISAFQQVLYYQNNVFMSESKCWVEHDINNGPHEFC 809
HCoV_HKU1 SMMILSDDGVVCYNSDYASKGYIANISVFQQVLYYQNNVFMSESKCWVENDITNGPHEFC 809
MHV SMMILSDDGVVCYNSEFASKGYIANISAFQQVLYYQNNVFMSEAKCWVETDIEKGPHEFC 809
MERS SMMILSDDGVVCYNSDYAAKGYIAGIQNFKETLYYQNNVFMSEAKCWVETDLKKGPHEFC 814
HKU4 SMMILSDDGVVCYNSDYAAKGYVASIQNFKETLYYQNNVFMSEAKCWVETNLEKGPHEFC 815
HKU5 SMMILSDDGVVCYNSDYATKGYIASIQNFKETLYYQNNVFMSEAKCWVETDLKKGPHEFC 815
HKU9 SMMILSDDGVVCYNADYAQKGYVADIQGFKELLYFQNNVFMSESKCWVEPDITKGPHEFC 813
SARS-CoV-2 SMMILSDDAVVCFNSTYASQGLVASIKNFKSVLYYQNNVFMSEAKCWTETDLTKGPHEFC 813
SARS-CoV SMMILSDDAVVCYNSNYAAQGLVASIKNFKAVLYYQNNVFMSEAKCWTETDLTKGPHEFC 813
.:******.*.* :    *  * ::.:  *:   *::**.*:*: :*** * ::   ******
Polymerase
Motif C
Motif D
Motif E
δ
α
γ
β
HKU19 SQHTVLAEYDGEPCYYPYPDVSRILGACIFVNETEKTDPVQNLERYISLAIDAYPLTKVD 880
HKU11 SQHTVLAEHEGKPYYLPYPDVSRILGACIFVDDVNKADPIQNLERYISLAIDAYPLTKVD 871
PDCV SQHTVLAEHDGKPYYLPYPDVSRILGACIFVDDVNKADPVQNLERYISLAIDAYPLTKVD 871
FCoV_65F SQHTLQIVGADGDYYLPYPDPSRILSAGVFVDDIVKTDNVIMLERYVSLAIDAYPLTKHP 870
HCoV_229E SQHTMQIVDENGKYYLPYPDPSRIISAGVFVDDVTKTDAVILLERYVSLAIDAYPLSKHP 868
HCoV_NL63 SQHTMQIVDKDGTYYLPYPDPSRILSAGVFVDDVVKTDAVVLLERYVSLAIDAYPLSKHP 868
HKU8 SQHTMQIVDGDGTYYLPYPDPSRILSAGVFVDDVIKTDAVVLLERYVSLAIDAYPLSKHP 868
PEDV SQHTMQIVDKDGTYYLPYPDPSRILSAGVFVDDVVKTDAVVLLERYVSLAIDAYPLSKHE 868
HKU22 SQHSMLVEIDGEMRYLPYPDPSRILGACVFVDDVEKTEPVVVMERYVALAIDAYPLIYHE 867
IBV SQHTMLVEVDGEPRYLPYPDPSRILCACVFVDDLDKTESVAVMERYIALAIDAYPLVHHE 881
HCoV_OC43 SQHTMLVKMDGDDVYLPYPNPSRILGAGCFVDDLLKTDSVLLIERFVSLAIDAYPLVYHE 869
HCoV_HKU1 SQHTMLVKIDGDYVYLPYPDPSRILGAGCFVDDLLKTDSVLLIERFVSLAIDAYPLVYHE 869
MHV SQHTMLVKMDGDEVYLPYPDPSRILGAGCFVDDLLKTDSVLLIERFVSLAIDAYPLVYHE 869
MERS SQHTLYIKDGDDGYFLPYPDPSRILSAGCFVDDIVKTDGTLMVERFVSLAIDAYPLTKHE 874
HKU4 SQHTLYIKDGDDGYFLPYPDPSRILSAGCFVDDIVKTDGTVMMERYVSLAIDAYPLTKHD 875
HKU5 SQHTLFIKDGDDGYFLPYPDPSRILSAGCFVDDIVKTDGTLMVERFVSLAIDAYPLTKHD 875
HKU9 SQHTMLVEMNGEKVYLPYPDPSRILGAGCFVDDLLKTDGTLMMERYVSLAIDAYPLTKHA 873
SARS-CoV-2 SQHTMLVKQGDDYVYLPYPDPSRILGAGCFVDDIVKTDGTLMIERFVSLAIDAYPLTKHP 873
SARS-CoV SQHTMLVKQGDDYVYLPYPDPSRILGAGCFVDDIVKTDGTLMIERFVSLAIDAYPLTKHP 873
***::      .   : ***: ***: *  **::  *::     :**:::********
Polymerase
Motif E
PEDV nsp12
V842
V848
SARS-CoV-2 nsp12
I847
T853

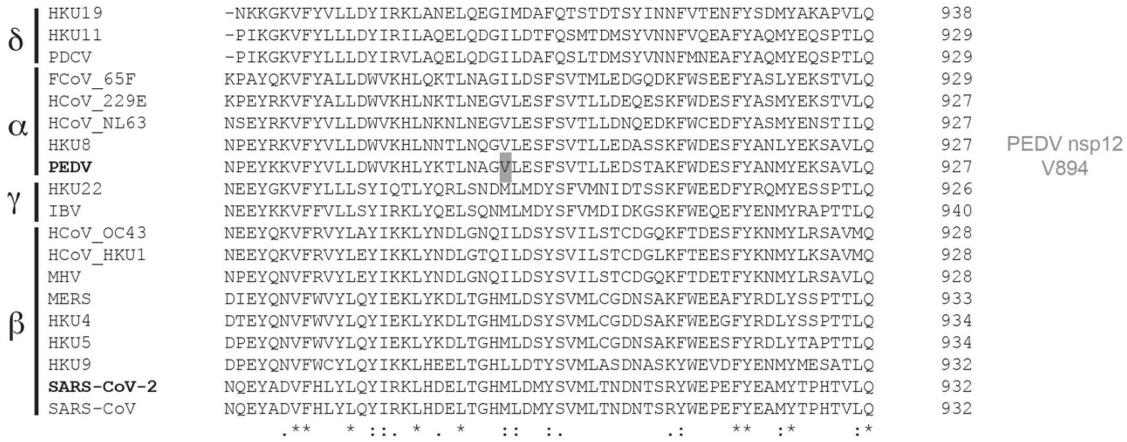


**Figure S5, Multiple sequence alignment of coronavirus nsp12s.** Alignment of nsp12 from the alpha- (FCoV-65F, HCoV-229E, HCoV-NL63, HKU8, PEDV), beta- (HCoV-OC43, HKU1, MHV MERS, HKU4, HKU5, HKU9, SARS-CoV, SARS-CoV-2), gamma (HKU22, IBV), and delta- (HKU19, HKU11, PDCV) coronavirus genera. Global alignment was done using Clustal Omega [2]. Residues marked with “*” are conserved, “:” are very similar, and “.” are moderately similar residues.

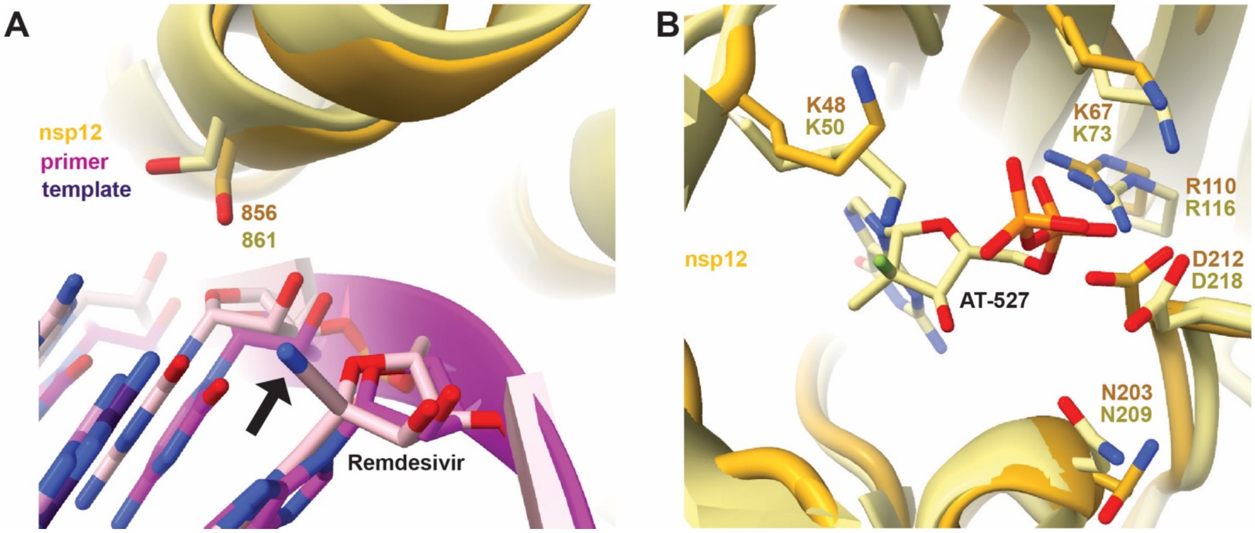


**Figure S6: possible cross-effectiveness of CoV antivirals.** In each figure PEDV nsp12 is shown in dark yellow and the RNA primer in dark pink, superimposed are SARS-CoV-2 models in lighter, matching colors. **A)** The antiviral Remdesivir's 1'-cyano group (black arrow) is believed to clash with SARS-CoV-2 nsp12 S861 in the +4 extension position. Remdesivir incorporated into a nascent primer at +3 is shown (PDB ID: 7B3C). Due to the sequence and spatial homology of S856 in PEDV we predict Remdesivir would have similar effectiveness against PEDV. **B**) The dual action antiviral AT-527 was shown to bind and inhibit the NiRAN domain of SARS-CoV-2 nsp12 (PDB ID: 7ED5). Several residues important for drug binding are conserved in PEDV, shown are K48, K67, R110, N203, and D212.

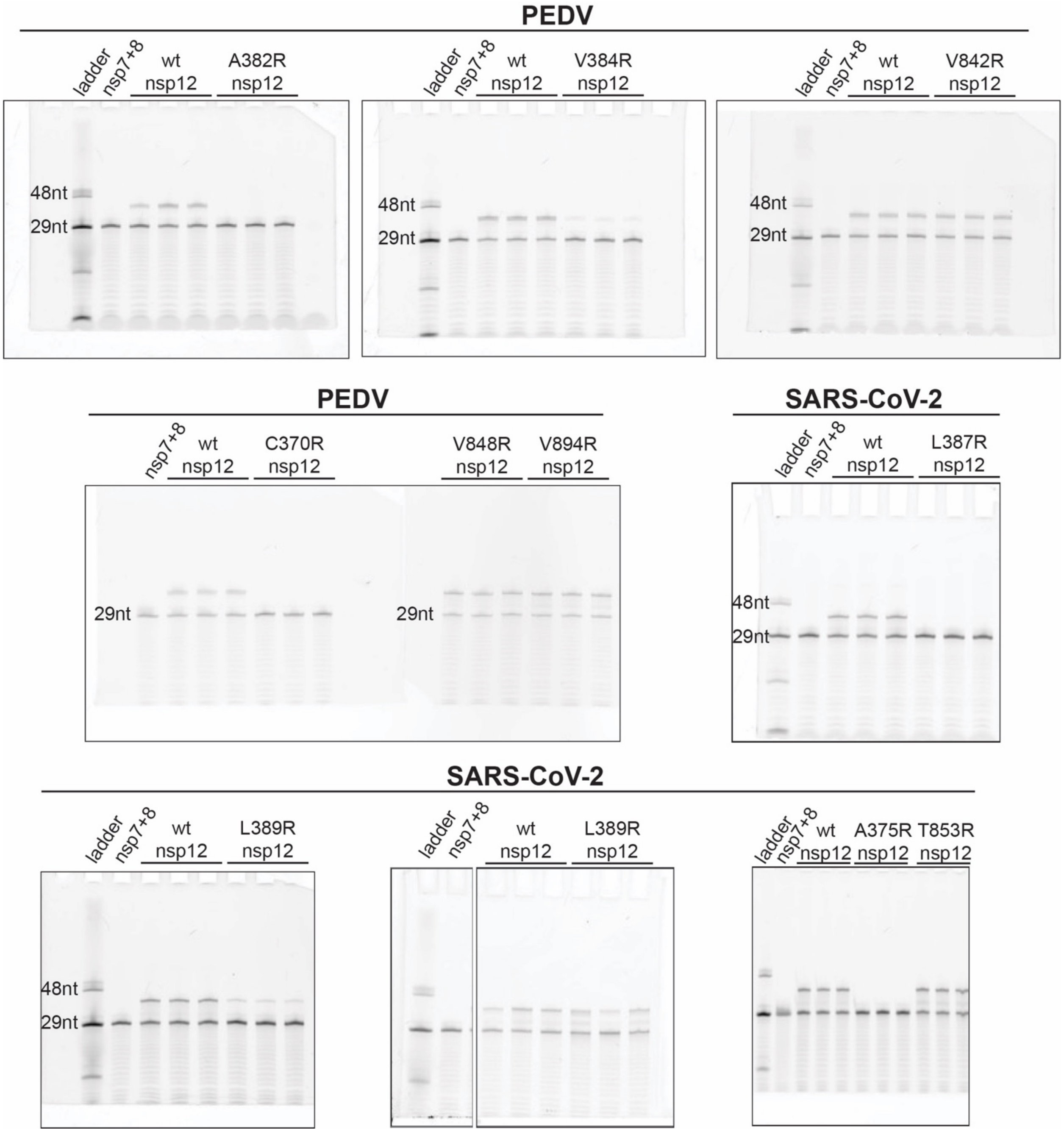


**Figure S7, full gel images for mutant nsp12 primer-extension assays:** Primer extension results for complexes with mutant nsp12s for PEDV and SARS-CoV-2. For each experiment wildtype (wt) and mutant polymerase reactions were done in triplicate. Viral cofactors, nsp7 and nsp8, were always provided in excess to nsp12 (wt or mutant). Each experiment included one negative control reaction lacking nsp12.

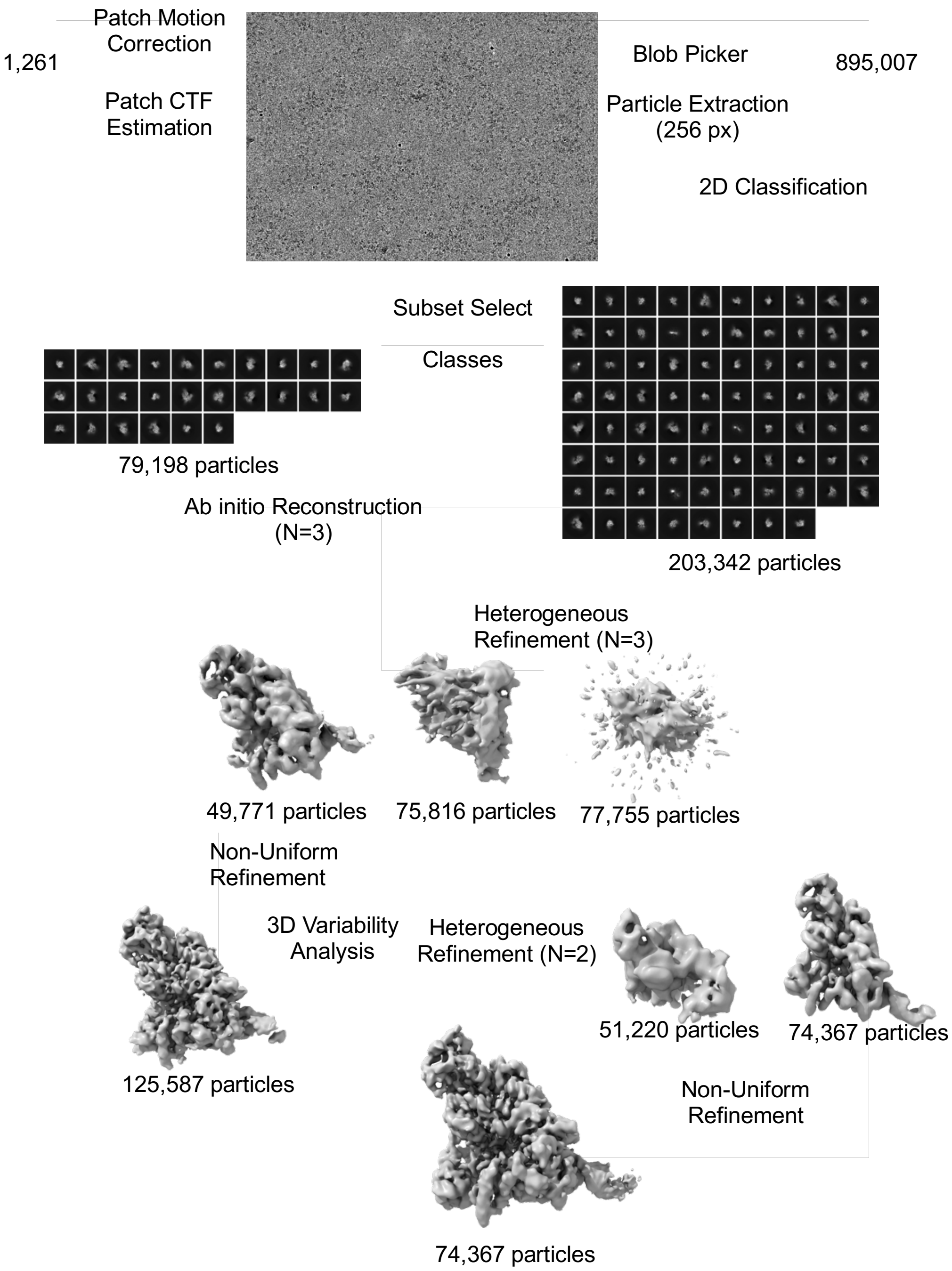


**Figure S8, cryo-EM processing pipeline:** Workflow for the processing of cryo-EM data starting with 1,261 movies. All data processing was performed in cryoSPARC [1]. Poor particles were removed through multiple rounds of 2D classification, then a subset of polymerase classes were used to generate initial models for subsequent heterogenous refinement with all remaining particles. Particles from classes resembling polymerases were merged for further 3D refinement and a final round of heterogenous refinement into nsp12 alone and polymerase complex maps. The complex map was further refined to our final 3.3 Å reconstruction.
